# ZATT/ZNF451 promotes release of stalled TOP2 cleavage complexes

**DOI:** 10.64898/2026.07.14.738426

**Authors:** Xueyuan Leng, Alessandra Zarantonello, Sampath Amitash Gadi, Ellen Kakulidis, Pierre Fey, Andreas Ingham, Ivo Hendriks, Sidak Minocha, Camilla Colding- Christensen, Stella M. Kristensen, Simon Willaume, Natalie Palkova, Christl Gaubitz, Alejandro García-López, Poul Martin Bendix, Claus Storgaard Sørensen, Michael Lund Nielsen, Norman E. Davey, Niels Mailand, Thomas C.R. Miller, Julien P. Duxin

**Author notes:** These authors contributed equally (Ellen Kakulidis, Pierre Fey, Andreas Ingham, Ivo Hendriks).

## Abstract

Topoisomerase II (TOP2) resolves DNA topological constraints through a tightly regulated cycle of DNA double-strand cleavage and religation. Nearby DNA damage or chemotherapeutic agents such as etoposide block the DNA religation step, stabilizing TOP2-DNA cleavage complexes (TOP2ccs) at DNA double-strand breaks (DSBs). The SUMO E3 ligase ZATT (ZNF451) has recently emerged as a key effector of TOP2cc repair, but its mechanism of action remains poorly understood. Here, we show that ZATT is sufficient to resolve TOP2ccs independently of TDP2, TOP2 proteolysis, and canonical DSB repair pathways. Using *Xenopus* egg extracts and biochemical reconstitution, we find that ZATT salvages trapped TOP2 by promoting TOP2 release from its stalled cleavage complex. Structural modeling and targeted mutagenesis in *Xenopus* egg extracts and human cells identify a highly conserved hydrophobic pocket in the tower domain of TOP2 where the ZATT coiled-coil “hooks on” to promote TOP2cc resolution. Our findings reveal a new strategy to resolve TOP2ccs that bypasses the exposure of dangerous DNA breaks.

## Introduction

DNA topoisomerases alleviate topological stress in the genome (Pommier et al. 2022). Among these, type II topoisomerases possess the unique ability to disentangle DNA knots and catenanes by passing one double helix through another via a highly regulated cycle of DNA double-strand breakage and religation (Nitiss 2009a). Vertebrates encode two type II topoisomerases: TOP2a and TOP2b (Tsai-Pflugfelder et al. 1988; Tan et al. 1992). Despite their high sequence homology, these enzymes carry out distinct functions (Nitiss 2009a; J. H. Lee and Berger 2019). TOP2a is essential for cell viability, playing critical roles in relieving torsional stress and resolving catenanes during DNA replication, as well as in chromosome condensation and segregation during mitosis (Holm et al. 1985; Uemura et al. 1987; Grue et al. 1998; Bermejo et al. 2007; Lucas et al. 2001). In contrast, TOP2b is primarily involved in transcriptional regulation and maintaining three-dimensional genome architecture (Lyu et al. 2006; Ju et al. 2006; Madabhushi et al. 2015; Pommier et al. 2022).

TOP2a and TOP2b function as homodimers and catalyze breakage and rejoining of duplex DNA in an ATP-dependent manner (**Fig.1A**) (Pommier et al. 2022). During catalysis, TOP2 introduces a transient DNA double-strand break (DSB) in one DNA duplex, known as the G-segment, to allow the passage of another duplex, termed the T-segment, through the enzyme’s DNA gate (**Fig.1A**, i-ii). The DSB results from staggered incisions four base pairs apart, formed via paired transesterification reactions, where each TOP2 subunit covalently links to the 5′ end of the cleaved DNA through a phosphotyrosyl bond, forming the TOP2-DNA cleavage complex (TOP2cc) (**Fig.1A**, ii). Prior to DNA incisions, ATP binding induces dimerization of the N-terminal ATPase domains, capturing the T-segment (**Fig.1A**, i) (Roca and Wang 1992). After DNA cleavage, hydrolysis of one ATP molecule is postulated to open the DNA gate, allowing T-segment passage (**Fig.1A**, ii) (Bjergbaek et al. 2000; Chen et al. 2018; Vanden Broeck et al. 2021). The DSB is then resealed, releasing the cleavage complex (**Fig.1A**, iii). Hydrolysis of the second ATP molecule reopens the N-gate and releases the T-segment from the C-gate, resetting the enzyme for another catalytic cycle (**Fig.1A**, iii–iv) (Schmidt, Osheroff, and Berger 2012; Ling et al. 2022).

**Figure 1.**
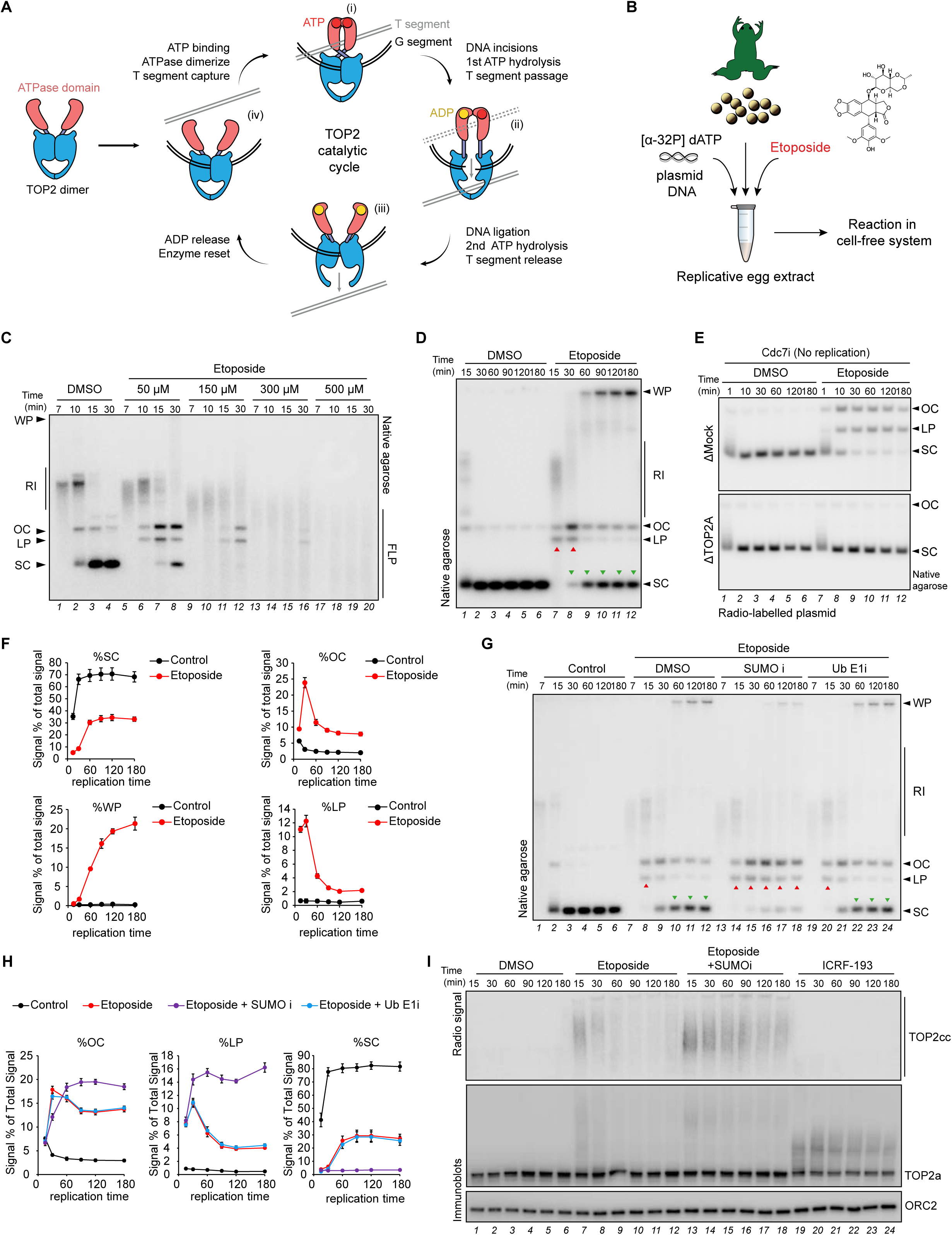
SUMOylation orchestrates TOP2cc resolution in Xenopus egg extracts. **A**) Schematic of the TOP2 catalytic cycle. **B**) Illustration of experimental set up in *Xenopus* egg extracts. **C**) pBluescript II SK(+) (pBS) was replicated in egg extracts supplemented with [α-^32^P] dATP, in the presence or absence of etoposide at indicated concentrations. At the indicated timepoints, reaction samples were treated with Proteinase K (ProtK) and analyzed by native gel electrophoresis. WP, well products; RI, replication intermediates; OC, open circular; LP, linear products; SC, supercoiled. FLP, fragmented linear products. **D**) pBS plasmid was replicated in egg extracts up to 180 min, supplemented with [α-^32^P] dATP, in the absence or presence of etoposide (100 µM). Reaction samples were analyzed as in (C). **E**) pBS was pre-radiolabeled *in vitro* with [α-^32^P] dATP and then was incubated in extract treated with etoposide. Cdc7 inhibitor (Cdc7i) was supplemented to block DNA replication. **F**) Quantification of SC, OC, WP and LP in (D), mean and standard error of three independent experiments is plotted. **G**). pBS plasmid was replicated in egg extracts in the absence or presence of etoposide. Where indicated, the SUMO– (ML792; SUMO i) or ubiquitin– (MLN-7243; Ub E1i) inhibitor was supplemented to the reaction. **H**) Quantification of OC, LP and SC in (G), mean and standard error of three independent experiments is plotted. **I**) pBS was replicated in the presence of DMSO, etoposide (100 µM), etoposide and SUMOi, or ICRF-193 (100 µM). Samples were analyzed via drag-on blot (radio signal) and immunoblot with indicated antibodies.

Although TOP2 is highly efficient, its catalytic cycle can be disrupted by nearby DNA lesions that block religation, leading to the stabilization of TOP2ccs. This block can also be induced by TOP2 poisons like etoposide, which intercalates at the DNA break and prevents religation (Wu et al. 2011). In contrast, other inhibitors, such as bisdioxopiperazines (e.g., ICRF-193 and ICRF-187), trap TOP2 through a noncovalent mechanism by locking the ATPase dimer interface and blocking the second ATP hydrolysis step, thereby preventing enzyme release after ligation (Classen, Olland, and Berger 2003; Ling et al. 2022).

TOP2-targeting agents like etoposide have been used clinically for over five decades to treat cancers including lung, testicular, ovarian and other tumors (Aisner and Lee 1991; Nitiss 2009b). Yet, the mechanisms by which TOP2ccs are repaired remain incompletely understood. Early studies showed that etoposide induces SUMOylation and ubiquitylation of TOP2, marking it for proteasomal degradation (Mao, Desai, and Liu 2000; Mao et al. 2001). A major advance came with the discovery of tyrosyl-DNA phosphodiesterase 2 (TDP2), which hydrolyzes the phosphotyrosyl bond between TOP2 and DNA (Cortes Ledesma et al. 2009). This underscored that TOP2ccs arise endogenously and are actively resolved by cells. However, TDP2 cannot hydrolyze full-length TOP2ccs *in vitro* as the crosslink is buried within the TOP2 catalytic core. This led to a consensus model in which proteasomal degradation of TOP2 precedes TDP2 activity, thereby granting the enzyme access to the phosphotyrosyl bond (Gao et al. 2014; Sun et al. 2020).

A key breakthrough in 2017 identified ZNF451, renamed ZATT (for **Z**inc finger protein **A**ssociated with **T**DP2 and **T**OP2), as a SUMO ligase that rapidly modifies TOP2ccs (Schellenberg et al. 2017). This work provided compelling molecular and structural evidence that ZATT-mediated SUMOylation facilitates the recruitment of TDP2 via a split SUMO-interacting domain (Schellenberg et al. 2017). Notably, this mechanism allows TDP2 to resolve TOP2ccs without requiring proteolytic degradation of TOP2.

Crucially, all known repair pathways for TOP2ccs converge on the generation of a toxic DSB intermediate. Once the protein-DNA crosslink is removed, whether by proteolysis or enzymatic cleavage, the resulting DSB must be repaired to maintain genomic integrity. The primary pathway for this repair is non-homologous end joining (NHEJ) (Gómez-Herreros et al. 2013). The MRN nuclease complex can also process TOP2ccs by incising near the crosslink, removing the protein obstruction and enabling DSB repair via NHEJ (Deshpande et al. 2016; Hoa et al. 2016; Aparicio et al. 2016; Hartsuiker, Neale, and Carr 2009; Quennet et al. 2011; Anand et al. 2016). More recently, the SWI2/SNF2-family helicases, Rrp2 in yeast and RAD54L2 in vertebrates, were shown to counteract etoposide cytoxicity, by displacing TOP2 from DNA (Wei et al. 2017; H. Zhang et al. 2023; D’Alessandro et al. 2023).

Based on cellular sensitivity to etoposide, ZATT and TDP2 appear to be the primary responders to TOP2ccs (Fig.S1A) (Olivieri et al. 2020). Notably, ZATT-deficient cells are more sensitive to etoposide than TDP2-deficient cells, and double knockouts show additive sensitivity (Schellenberg et al. 2017; Park et al. 2023), suggesting ZATT has TDP2-independent functions in TOP2cc resolution. Further highlighting their distinct roles, TDP2 knockout cells are sensitive to the non-covalent TOP2 trapper ICRF-187, whereas ZATT knockouts are not (Fig.S1B) (Olivieri et al. 2020). The nature of ZATT’s TDP2-independent functions remains unknown and represents a key unresolved question in the field.

Here, we recapitulate TOP2cc repair in *Xenopus* egg extracts and show that ZATT orchestrates TOP2cc repair in this system. Remarkably, ZATT promotes TOP2cc resolution independently of TDP2, TOP2 proteolysis, or canonical DSB repair pathways. A CRISPR tiling screen in human cells identifies key regions of ZATT required for this activity, and *in vitro* reconstitution shows that ZATT enables TOP2 release from its trapped cleavage complex. We identify a conserved binding pocket on TOP2 essential for ZATT– mediated repair. Collectively, these findings uncover a safe and efficient mechanism of TOP2cc repair mediated by ZATT, which bypasses proteolytic degradation and avoids the exposure of hazardous DSBs. This breakage-independent TOP2cc repair mechanism explains the exquisite sensitivity of ZATT-deficient cells to etoposide and highlights novel therapeutic opportunities to sensitize cancer cells to etoposide.

## Results

### Etoposide induces TOP2ccs in *Xenopus* egg extracts

To study TOP2cc repair in *Xenopus* egg extracts, we titrated etoposide during replication of an undamaged plasmid (**Fig.1B**). As expected, etoposide inhibited DNA replication in a dose-dependent manner (**Fig.1C**) (Van Ravenstein et al. 2022). When both subunits of one TOP2 dimer are simultaneously crosslinked to a plasmid, a linear DNA product (LP) appears after proteinase K treatment, whereas crosslinking of only one of the two subunits yields nicked open circular (OC) DNA molecules. Low dose etoposide (∼50-100 µM) produced both LP and OC DNA species (**Fig.1C**, lanes 5-8, **Fig.1D**, lanes 7-12). Higher doses generated a smear of fragmented linear products (FLPs), suggesting that multiple TOP2 dimers were crosslinked to the same plasmid (**Fig.1C**, lanes 9-20). In contrast, treatment with ICRF-193 led to catenated DNA (Fig. S1C, Cats), consistent with inhibition of TOP2 catalytic activity through non-covalent trapping (J. H. Lee, Wendorff, and Berger 2017). Etoposide-induced LP and OC products also formed without replication and were largely dependent on TOP2a (**Fig.1E**, S1D). We conclude that etoposide efficiently crosslinks TOP2a to plasmid DNA in egg extracts with no apparent contribution from TOP2b.

Treatment with 100 µM etoposide also led to the appearance of well products (WPs) between 60-180 minutes, likely derived from processed aberrant replication intermediates (**Fig.1D**, lanes 9-12, **Fig.1F**) (Van Ravenstein et al. 2022). WP formation required active DNA replication (Fig.S1E) and was observed when etoposide was added during replication (11 min), but not after its completion (31 min) (Fig.S1F). This suggests that WPs arise from replication forks encountering TOP2ccs, possibly through fork reversal and processing (Tian et al. 2021). WP generation was also dependent on TOP2a (Fig.S1G).

LP and OC species, which were predominant at 15– and 30-minutes following decatenation (**Fig.1D**, lanes 7-8; **Fig.1F**), decreased in intensity at 60 to 180 minutes (**Fig.1D**, lanes 9-12; **Fig.1F**). This decline coincided with an increase in supercoiled (SC) plasmids, suggesting that the initial LP and OC molecules are repaired and converted to SC forms. Since etoposide remains present, the reaction likely reaches a dynamic equilibrium around 60 minutes between TOP2cc formation and repair. Consequently, this concentration of etoposide provided an optimal window to monitor TOP2cc formation and resolution.

Given the established roles of SUMOylation and ubiquitylation in DNA-protein crosslink resolution and their implication in TOP2cc repair (Riccio, Schellenberg, and Williams 2020; Leng and Duxin 2022), we first tested the effect of inhibiting these signaling pathways. Reactions were supplemented with either a SUMO or ubiquitin E1-activating enzyme inhibitor, which we previously shown to work in egg extract (He et al. 2017; Hyer et al. 2018; Liu et al. 2021; Gallina et al. 2021). Inhibition of SUMOylation reduced fully repaired SC products and led to accumulation of both OC and LP species (**Fig.1G**, lanes 13-18, **Fig.1H**). In contrast, inhibition of ubiquitylation had no appreciable effect on the reaction (**Fig.1G**, lanes 19-24, **Fig.1H**). These results indicate that SUMOylation, but not ubiquitylation, plays a critical role in regulating the generation or repair of TOP2ccs in egg extracts. Notably, SUMOylation also regulated TOP2cc resolution independently of DNA replication (Fig.S1H).

To monitor TOP2cc formation and resolution, we developed the “drag-on blot” assay, which detects radiolabeled DNA covalently linked to TOP2a following SDS-PAGE and transfer to a PVDF membrane (Fig.S1I). To validate the specificity of the assay, we depleted TOP2a during DNA replication to accumulate plasmid catenanes, then supplemented extracts with either wild-type TOP2a (TOP2a WT) or catalytically inactive mutant TOP2a (TOP2a CD) and assessed decatenation and TOP2cc formation. TOP2a WT, but not TOP2a CD, efficiently decatenated plasmids (Fig.S1K-L) and transferred radiolabeled DNA to the membrane in the presence of etoposide (Fig.S1M). These results confirm that the drag-on blot specifically detects covalent TOP2ccs and can be used to track their formation and resolution in egg extracts. Notably, etoposide treatment produced a drag-on blot signal, which was enhanced and sustained upon SUMO inhibition, consistent with impaired repair (**Fig.1I**, lanes 7-18). In contrast, treatment with ICRF-193, which traps TOP2a in a non-covalent manner, did not produce a radio signal (**Fig.1I**, lanes 19-24). Collectively, these results indicate that in *Xenopus* egg extracts the repair of etoposide-induced TOP2cc is regulated by SUMOylation but not ubiquitylation.

### TOP2cc resolution is governed by ZATT in egg extracts

To investigate how TOP2cc repair occurs in *Xenopus* egg extracts, we performed plasmid pull-down mass spectrometry (PP-MS) to identify proteins enriched on replicating plasmids in the presence of etoposide with and without inhibition of SUMOylation (**Fig.2A**) (Larsen et al. 2019). As shown in **Fig.2B**, replisome components persisted on the plasmid treated with etoposide for up to 30 minutes, consistent with a delay in completing replication of the plasmid (**Fig.2A**, compare lanes 2 and 6). Additionally, we observed the accumulation of replication fork reversal and homologous recombination (HR) factors, implicating these pathways in processing stalled replication forks (**Fig.2B**, Table S1) (Adolph and Cortez 2024).

**Figure 2.**
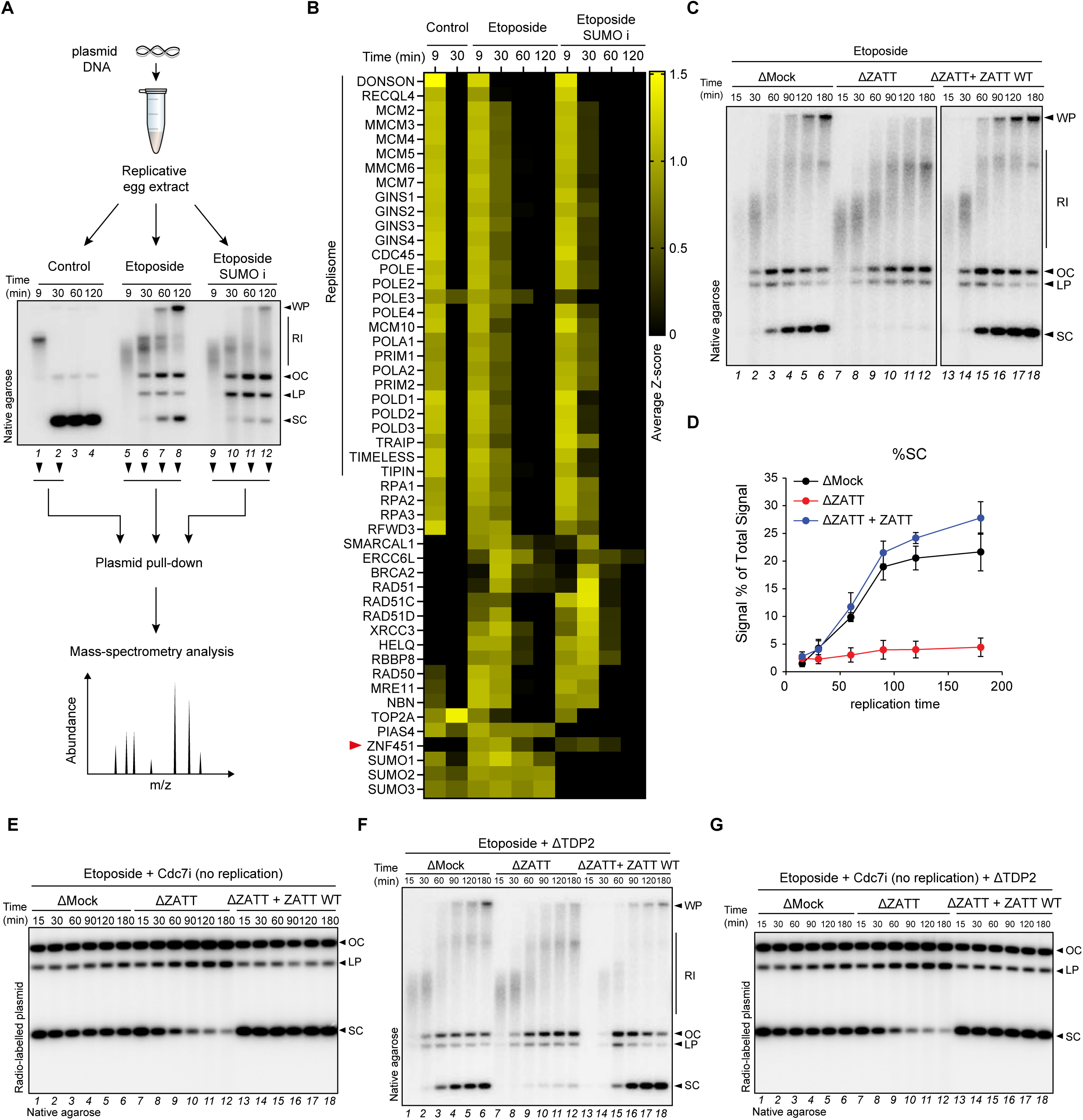
ZATT governs TOP2cc resolution independently of TDP2. **A**) Workflow illustration of plasmid pull-down mass spectrometry (PP-MS). pBS was replicated in the presence of DMSO, etoposide (100 µM), or etoposide and SUMO i. Reactions were analyzed as in Fig.1C. Four independent reactions from indicated timepoints were used for PP-MS (Fig.2B). **B**) Heatmap of PP-MS analysis displaying the mean of the z-scored Log_2_ label-free quantification (LFQ) intensity for the indicated proteins obtained from four independent replicates of pBS replication (Fig.2A). Either etoposide or etoposide and SUMOi were supplemented where indicated. **C**) pBS was replicated in the presence of etoposide in mock or ZATT-depleted egg extracts. ZATT-depleted extracts were supplemented with recombinant ZATT wildtype (WT) where indicated. Reaction samples were analyzed as in Fig.1C. **D**) Quantification of SC product in (C), mean and standard error of three independent experiments is plotted. Differently lettered groups are significantly different from each other, by one-way ANOVA with Tukey HSD test. **E**) pBS was pre-radiolabeled *in vitro* with [α-^32^P] dATP and then incubated in extracts from (C) in the presence of etoposide and Cdc7i to block DNA replication initiation. Reaction samples were analyzed as in Fig.1E. Note the generation of fragmented linear products (FLP) upon ZATT depletion (lanes 10-12). **F**) Same as (C) but TDP2 were depleted from extract. **G**) Same as (E) but extracts were depleted of TDP2.

When monitoring the SUMO ligases known to participate in TOP2cc repair, we detected the recruitment of both ZATT (ZNF451) and PIAS4 (**Fig.2B**, S2A, Table S1). While PIAS4 was present on the replicating plasmid independently of etoposide, ZATT recruitment was specific to the etoposide treatment (**Fig.2B**, S2A). Notably, we did not detect TDP2 recruitment, although TDP2 is present in egg extracts (Colding-Christensen et al. 2023).

Given that ZATT deficiency confers the highest sensitivity to etoposide (**Fig.S1A**) (Schellenberg et al. 2017; Olivieri et al. 2020), we generated a specific antibody to deplete ZATT from extracts (Fig.S2B). Notably, depletion of ZATT impaired the formation of SC products both in the presence and absence of DNA replication when etoposide was supplemented to the reaction (**Fig.2C**-**E**, lanes 7-12). Addition of recombinant ZATT rescued this effect, restoring SC products (**Fig.2C**-**E**, lanes 13-18, S2C). Taken together, these findings show that *Xenopus* egg extracts recapitulate ZATT-dependent TOP2cc repair, providing an opportunity to dissect the molecular mechanisms underlying this process.

### ZATT-mediated TOP2cc repair occurs independently of TDP2

Given the absence of TDP2 recruitment to plasmids treated with etoposide (**Fig.2B**, Table S1), we investigated whether ZATT-dependent TOP2cc repair occurs independently of TDP2. To this end, we generated a TDP2 antibody that efficiently depletes the protein from extracts (Fig.S2D-E, Table S2). As shown in Fig.S2F, TDP2 depletion had no appreciable effect on the formation of SC products, suggesting that TDP2 is dispensable for TOP2cc repair in egg extracts. To confirm that ZATT acts independently of TDP2, we performed ZATT depletion and rescue in TDP2-depleted extracts. In this setting, ZATT depletion abolished repair both with and without replication (**Fig.2F-G**, lanes 7-12). Addback of recombinant ZATT fully restored SC product formation (**Fig.2F-G**, lanes 13– 18). Moreover, MS analysis of TOP2a or ZATT immunoprecipitates from egg extracts treated with etoposide revealed co-enrichment of ZATT and TOP2a, but not of TDP2 (Fig.S2G, Table S3). Although the lack of TDP2 recruitment to TOP2ccs in egg extracts remains unexplained, it may reflect a germline-specific regulation. Nonetheless, our data clearly show that ZATT can function in TOP2cc repair independently of TDP2, consistent with genetic observations in human cells (Park et al. 2023).

### TOP2cc repair occurs without proteolysis and DSB repair

Given that TOP2ccs can be targeted for proteasomal degradation (Riccio, Schellenberg, and Williams 2020), we next addressed whether the proteasome is involved in the repair of TOP2ccs. However, inhibition of de novo ubiquitylation or proteasome had little to no effect on the generation of SC products (**Fig.3A**, lanes 13-24, **Fig.3B**, Fig.S3A) or the accumulation of radio signal via dragon blot (**Fig.3C-D**) suggesting that TOP2cc proteolysis does not contribute to the ZATT pathway. TOP2cc proteolysis and/or removal leads to the exposure of DSBs, which become substrates of NHEJ (Riccio, Schellenberg, and Williams 2020). Thus, we examined the involvement of DSB repair factors in the pathway orchestrated by ZATT. To this end, we depleted XRCC4 (NHEJ), BRCA2 or RAD51 (HR) and replicated plasmids in the presence of etoposide (Fig.S3B). Unexpectedly, XRCC4, RAD51 or BRCA2 depletion did not hinder the generation of repaired SC products (**Fig.3E**). Similarly, inhibiting or depleting the microhomology-mediated end-joining factor Polθ did not show detectable effect of TOP2cc repair (Fig.S3B-C). Together, these results suggest that ZATT-mediated TOP2cc repair operates independently of canonical DSB repair factors.

**Figure 3.**
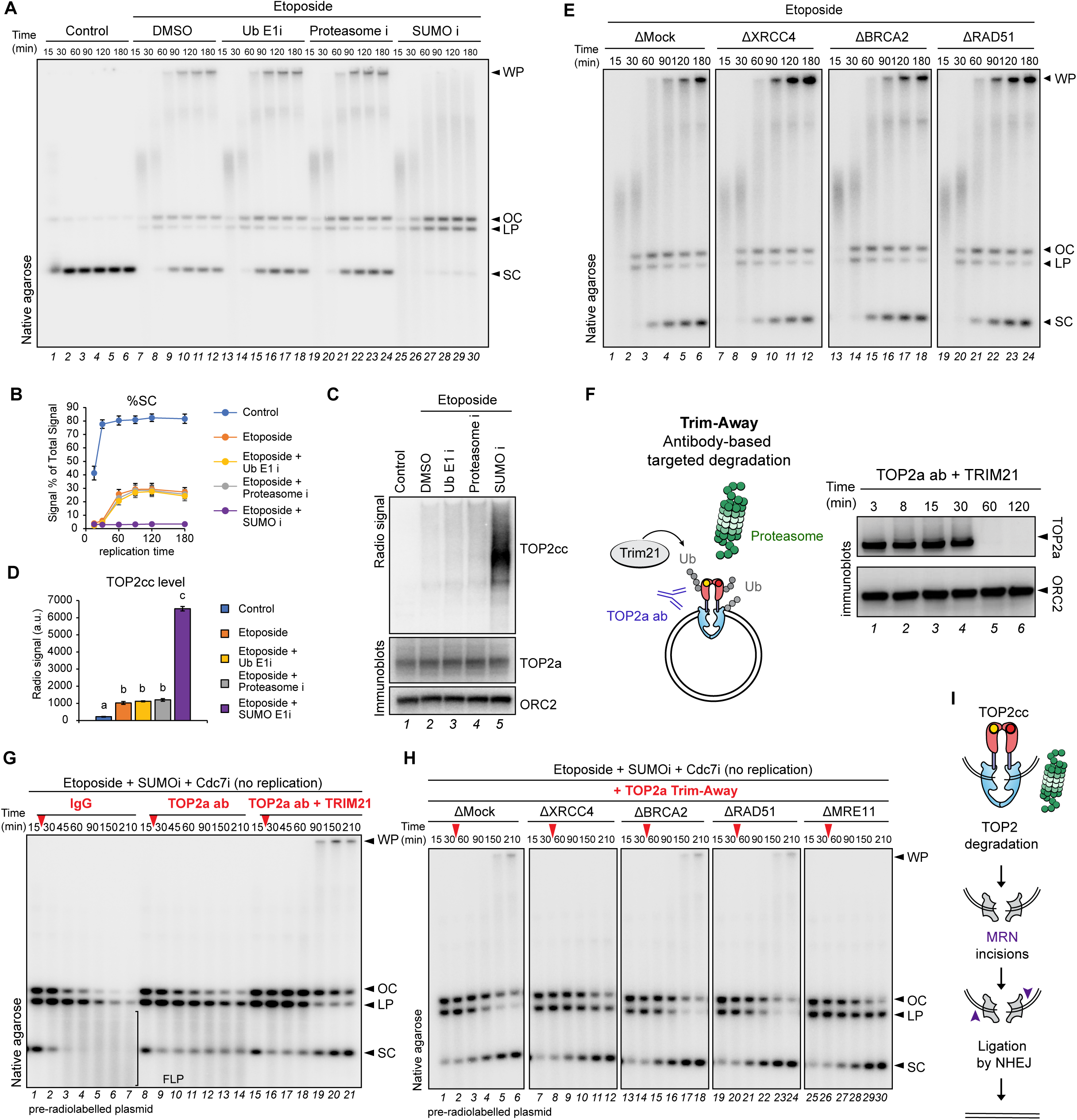
ZATT-mediated TOP2cc resolution is independent of the proteasome and DSB repair. **A**) pBS was replicated in egg extracts in the presence of etoposide (100 µM). DMSO, Ub E1i (MLN7243), proteasome inhibitor (MG262) or SUMOylation inhibitor (SUMO i) as indicated. Samples were analyzed as in Fig.1C. **B**) Quantification of SC in (A), mean and standard error of three independent experiments is plotted. **C**) Samples at 180 min from (A) were analyzed via drag-on blot (radio signal) and immunoblot with indicated antibodies. **D**) Quantification of radio-signal from (C), mean and standard error of three independent experiments is plotted. **E**) pBS was replicated in the presence of etoposide (100 µM) in either Mock-, XRCC4-, BRCA2-, or RAD51-depleted extracts. Samples were analyzed as in Fig.1C. **F**) Schematic of Trim-Away assay (left scheme). Recombinant TRIM21 and TOP2a antibody were used to induce targeted TOP2a degradation. Reaction samples of the indicated time points were immunoblotted with the indicated antibodies to assess TOP2a degradation (right panel). **G**) Pre-radiolabeled pBS was incubated in egg extracts supplemented with etoposide, SUMOylation inhibitor and Cdc7 inhibitor. At 15 minutes, either IgG (control), TOP2a antibody (control), or a mixture of TOP2a antibody and TRIM21 were added to the reaction (red arrow). Samples were analyzed as in Fig.1E. Note that TOP2a antibody addition appears to block TOP2a engagement with DNA leading to the stabilization of SC products. Resolution of FLPs is specific to the TOP2a antibody and TRIM21 condition. **H**) Pre-radiolabeled pBS was incubated in egg extracts either mock-, XRCC4-, BRCA2-, RAD51-, or MRE11-depleted. Reactions were supplemented with etoposide, SUMOylation inhibitor and Cdc7 inhibitor, and spiked with a mixture of TOP2a antibody and TRIM21 at 30 minutes (red arrow). Samples were analyzed as in Fig.1E. **I**) Illustration of TOP2cc repair following TOP2 proteolysis. DSBs are exposed and become a substrate for MRN and NHEJ.

On the basis of these results, we hypothesized that DSB repair factors might become necessary if TOP2ccs undergo proteolysis. To test this, we employed Trim-Away, an antibody-based method for targeted protein degradation via the proteasome (**Fig.3F**) (Clift et al. 2018; Mevissen, Prasad, and Walter 2023). By simultaneously adding the ubiquitin ligase TRIM21 and TOP2a antibodies to the extract, we achieved full TOP2a degradation by the proteasome within 30 to 60 minutes (**Fig.3F** and Fig.S3D). We next applied Trim-Away in extracts treated with etoposide. To facilitate the analysis, we performed these experiments in the absence of DNA replication and TOP2cc repair via SUMOi. Remarkably, while etoposide and SUMOi treatment led to the appearance of FLPs, induction of TOP2cc degradation via Trim-Away, restored the formation of repaired SC products with concomitant disappearance of FLPs (**Fig.3G**, compare lanes 1-7 and 15-21). TOP2cc degradation also resulted in the appearance of WPs, which could arise from the ligation of multiple linear molecules via NHEJ (**Fig.3G**, lanes 19-21). Consistent with this interpretation, the linear fragments were now stabilized in the absence of XRCC4, and WP never appeared (**Fig.3H**, compare lanes 1-6 and 7-12). In contrast, depletion of RAD51 or BRCA2 had no effect on the reaction (**Fig.3H**, lanes 13-18 and 19– 24). Given that the MRN endonuclease cleaves off proteins linked to the 5’ end of DNA (Reginato and Cejka 2020), we also generated an antibody to immunodeplete the MRN nuclease subunit MRE11 from egg extracts (Fig.S3B). Depletion of MRE11 also led to a strong stabilization of LPs and complete loss of WPs (**Fig.3H**, lanes 25-30). Based on these experiments, we conclude that TOP2ccs become substrates for MRN and NHEJ following TOP2 proteolysis (**Fig.3I**). In contrast, the ZATT-mediated resolution of TOP2ccs observed in egg extracts occurs independently of both proteolysis and DSB repair.

### ZATT SUMO ligase activity is not essential for TOP2cc repair

To systematically dissect ZATT domain functions in TOP2cc repair, we performed a CRISPR-Cas9 base editor screen using a single guide RNA (sgRNA) library targeting ZATT with missense mutations. The library was cloned into an ABE8e-SpG48 construct, enabling A-to-G conversions in a window overlapping the sgRNA target region (Sangree et al. 2022) and predicted to mutate >55% of ZATT’s amino acids (Table S4). RPE1– hTERT p53-/– cells were transduced with the library and cultured with etoposide, ICRF– 193, or left untreated for 12 days. sgRNA abundance was then quantified by next– generation sequencing to identify mutations sensitizing cells to drug treatment (**Fig.4A**).

**Figure 4.**
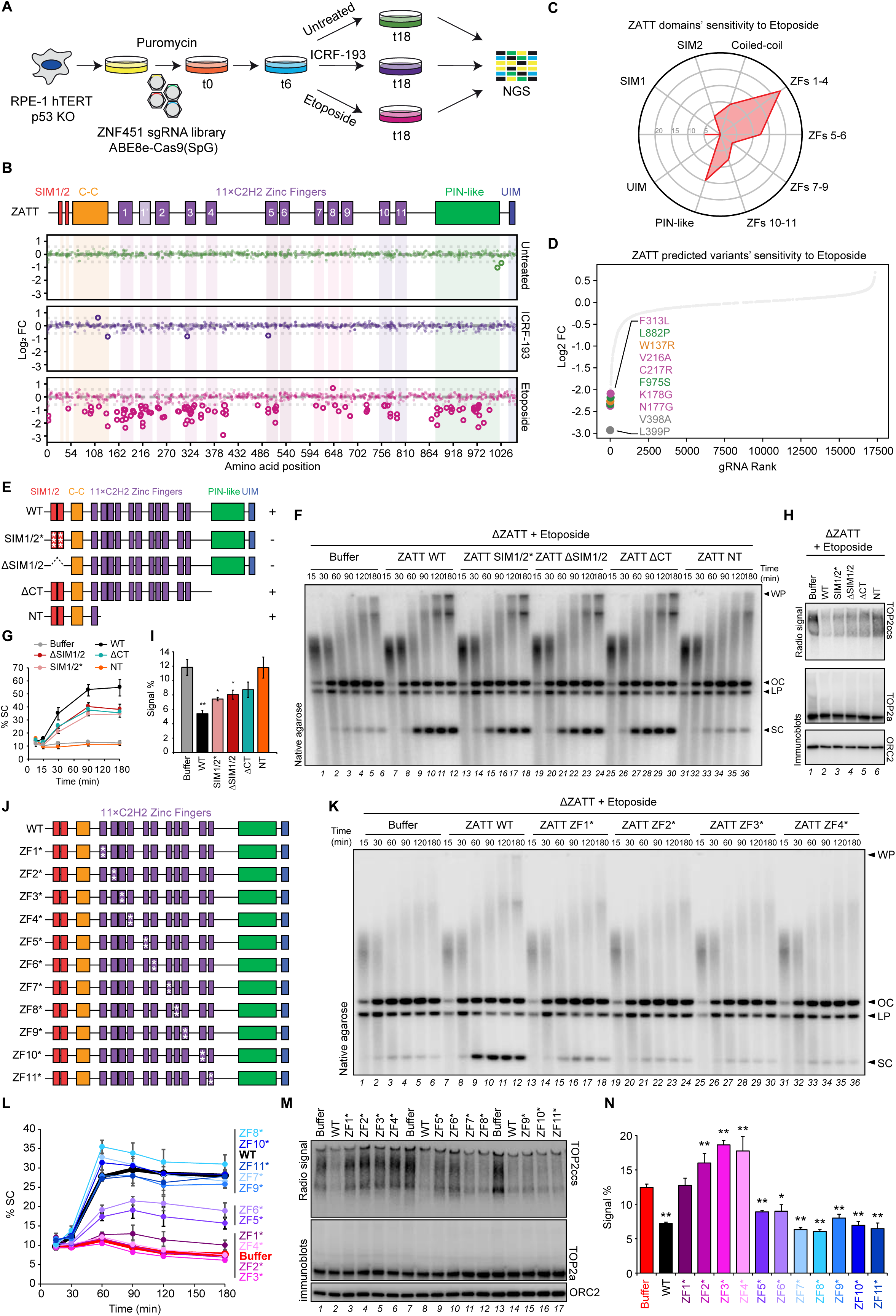
ZATT-mediated TOP2cc repair requires ZF1-4. **A**) Schematic outline of a ZATT CRISPR base editor tiling screen aimed at identifying missense mutations that sensitize RPE1-hTERT p53-/– cells to etoposide. Following selection (t6), cells were either left untreated or treated with a low dose ICRF-193 or etoposide for twelve additional days (t18). NGS, next generation sequencing. **B**) Dot plot showing the results of the ZATT CRISPR base editor tiling screen. Each guide is shown as a dot. X-axis represents the amino acid position in ZATT targeted for point mutation. Y-axis represents Log2 fold changes between ICRF-193– or etoposide– and untreated conditions. Larger hollow dots represent guides significantly changing (p-value ≤ 0.01). The various domains of ZATT are indicated above the dot blot. SIM, SUMO-interacting motif; C-C, coiled-coil; PIN-like, PilT N-terminus like domain; UIM, Ubiquitin-interacting motif. **C**) Spider plot illustrating the percentage of targeting guides that sensitize cells to etoposide in each group of indicated domains. **D**) Plot illustrating the predicted mutations for the 10 most etoposide-sensitizing guide RNAs. Note that V398A and L399P are in a predicted folded region of the protein linked to ZF4. **E**) Overview of *Xenopus* ZATT and the specific mutants used in (F-I). + and – denote SUMO ligase active and dead, as was assessed in Fig.S4B. **F**) pBS was replicated in ZATT-depleted extracts reconstituted with either buffer or the indicated mutants. Samples at the indicated time points were analyzed as in Fig.1C. **G**) Quantification of repaired SC products in (F). Mean and standard error from three independent experiments is plotted. **H**) Samples at 180 min timepoint from (F) were analyzed via drag-on blot (radio signal) and immunoblot with indicated antibodies. **I**) Quantification of TOP2ccs via drag-on blot at 180 minutes from (H). Mean and standard error of the mean (SEM) of three independent experiments are shown. * and ** denote p– values ≤ 0.05 and p-values ≤0.01, respectively, by one-way ANOVA with Dunnet test compared to the “Buffer” condition. **J**) Overview of *Xenopus* ZATT zinc finger (ZF) mutants. For each individual ZF, the signature C2H2 sequence was mutated to A2H2. **K**) pBS was replicated in ZATT-depleted extracts reconstituted with either buffer or the indicated ZATT mutants. Samples at the indicated time points were analyzed as in Fig.1C. **L**) Quantification of repaired SC products in (K) and Fig.S4D-E. Mean and standard error of three independent experiments is plotted. **M**) Samples at 180 min from (K) and Fig.S4D-E were analyzed via drag-on blot (radio signal) and immunoblot with indicated antibodies. **N**) Quantification of TOP2ccs via dragon blot from (M). Mean and standard error of the mean (SEM) of three independent experiments are shown. * and ** denote p– values ≤ 0.01 and p-values ≤0.05, respectively, by one-way ANOVA with Dunnet test compared to the “Buffer” condition.

ZATT is a multifunctional protein composed of distinct domains (**Fig.4B**). Its N-terminus contains SUMO-interacting motifs (SIM1/2) essential for SUMO ligase activity (Eisenhardt et al. 2015; Cappadocia, Pichler, and Lima 2015), followed by a coiled-coil domain and 12 zinc fingers. Of these, 11 are classical C2H2 types and one is an atypical variant (designated as ZFs 1-11 and 1′, respectively) (Riccio, Schellenberg, and Williams 2020). Toward the C-terminus, ZATT harbors a PIN-like domain and a ubiquitin-interacting motif (UIM).

Consistent with ZATT’s role in modulating etoposide sensitivity, we observed significant sensitization of targeting guides in the etoposide treatment condition (**Fig.4B**, Table S4). Most functional domains contributed to this effect, with the strongest impact from ZF1-4, the coiled-coil, and PIN-like domains (**Fig.4B-C**, Table S4). ZF5 also scored highly, whereas ZF7-11 showed comparatively weaker effects (**Fig.4B-C**). Among the ten most sensitizing gRNAs, seven targeted the region encompassing ZF1-4, two the PIN– like domain, and one the end of the coiled-coil domain (**Fig.4D**). Surprisingly, guides targeting the SUMO-interacting motifs (SIM1-2) did not confer sensitivity, suggesting that ZATT’s role in the etoposide response may be independent of its SUMO ligase activity (**Fig.4B**, Table S4). In contrast to etoposide, ICRF-193 treatment did not confer marked sensitivity, in accordance with previous findings (**Fig.4B**, Table S4) (Olivieri et al. 2020).

Given the lack of sensitization of ZATT SIM domains, we tested previously characterized SIM mutants (SIM1/2* and ΔSIM1/2) (Eisenhardt et al. 2015) in *Xenopus* egg extracts and confirmed their loss of SUMO ligase activity (**Fig.4E** and Fig.S4A-B). We also generated an N-terminal truncation (NT) that retains SUMO ligase function (Eisenhardt et al. 2015), as well as a C-terminal truncated mutant (ΔCT) lacking both the PIN-like and UIM domains (**Fig.4E** and Fig.S4A-B). Despite the importance of SUMO-signaling in resolving TOP2ccs, ZATT SIM mutants significantly rescued the repair defects caused by ZATT depletion, though less effectively than wild-type ZATT (ZATT WT) (**Fig.4F**, lanes 13-24, **Fig.4G-I**). In contrast, the NT construct, which retains SUMO ligase activity, failed to rescue any defect (**Fig.4F**, lanes 31-36, **Fig.4G-I**). We also observed substantial rescue with the ΔCT mutant, suggesting only a modest contribution of the PIN-like and UIM domains to ZATT’s function in egg extracts (**Fig.4F**, lanes 25-30, **Fig.4G-I**). These results indicate that ZATT’s SUMO ligase activity plays a minor role in TOP2cc repair, despite global SUMOylation inhibition blocking repair (**Fig.1G**). This discrepancy may reflect redundancy with other SUMO E3 ligases known to SUMOylate TOP2, such as PIAS4 (Ryu et al. 2010).

### ZF1-4 are critical for ZATT function in TOP2cc repair

Our tiling screen revealed a pattern of functional importance across ZATT’s zinc fingers: ZF1-4 scored strongly, ZF5-6 showed intermediate effects, while ZF7-11 scored poorly (**Fig.4B**). To investigate this further, we generated point mutants for each of the 11 typical C2H2 ZFs to selectively disrupt them (**Fig.4J**, Fig.S4C) and assessed their ability to support TOP2cc repair using replication assays and drag-on blot analysis. We found that among all the ZF disruptions, mutants of ZF1, ZF2, ZF3 or ZF4 failed to rescue the repair defect caused by ZATT depletion, confirming their critical role (**Fig.4K-N,** Fig.S4D-E). ZF5 and 6 mutants showed partial rescue compared to ZATT WT, consistent with their intermediate score in the tiling screen (**Fig.4K-N**, Fig.S4D-E). In contrast, mutants of ZF7, ZF8, ZF9, ZF10, and ZF11 rescued as efficiently as wild-type ZATT (**Fig.4K-N,** Fig.S4D– E). From these results, we conclude that ZFs 1-4 are essential for ZATT-mediated TOP2cc repair, while ZFs 5-11 play more supportive or redundant roles.

### ZATT stimulates TOP2cc repair

We hypothesized that ZATT may resolve TOP2ccs either by limiting TOP2a activity, preventing formation of TOP2ccs, or by directly promoting repair. Consistent with a recent report, we observed a very mild inhibition of TOP2a relaxation in vitro when ZATT and TOP2a were added at equimolar concentrations (Fig.S5A, lanes 33-38) (Terrón-Bautista et al. 2025). To test if ZATT inhibits TOP2a in the extract, TOP2a-depleted extracts were spiked in with recombinant TOP2a either alone or in combination with recombinant ZATT added up to 2-times the concentration required for ZATT depletion rescue (**Fig.5A**). Notably, we observed no inhibition of TOP2A-mediated decatenation. In contrast, ZATT addition enhanced SC product formation compared to control in the presence of etoposide, suggesting that it promotes TOP2cc resolution rather than prevents their formation (**Fig.5B**).

**Figure 5.**
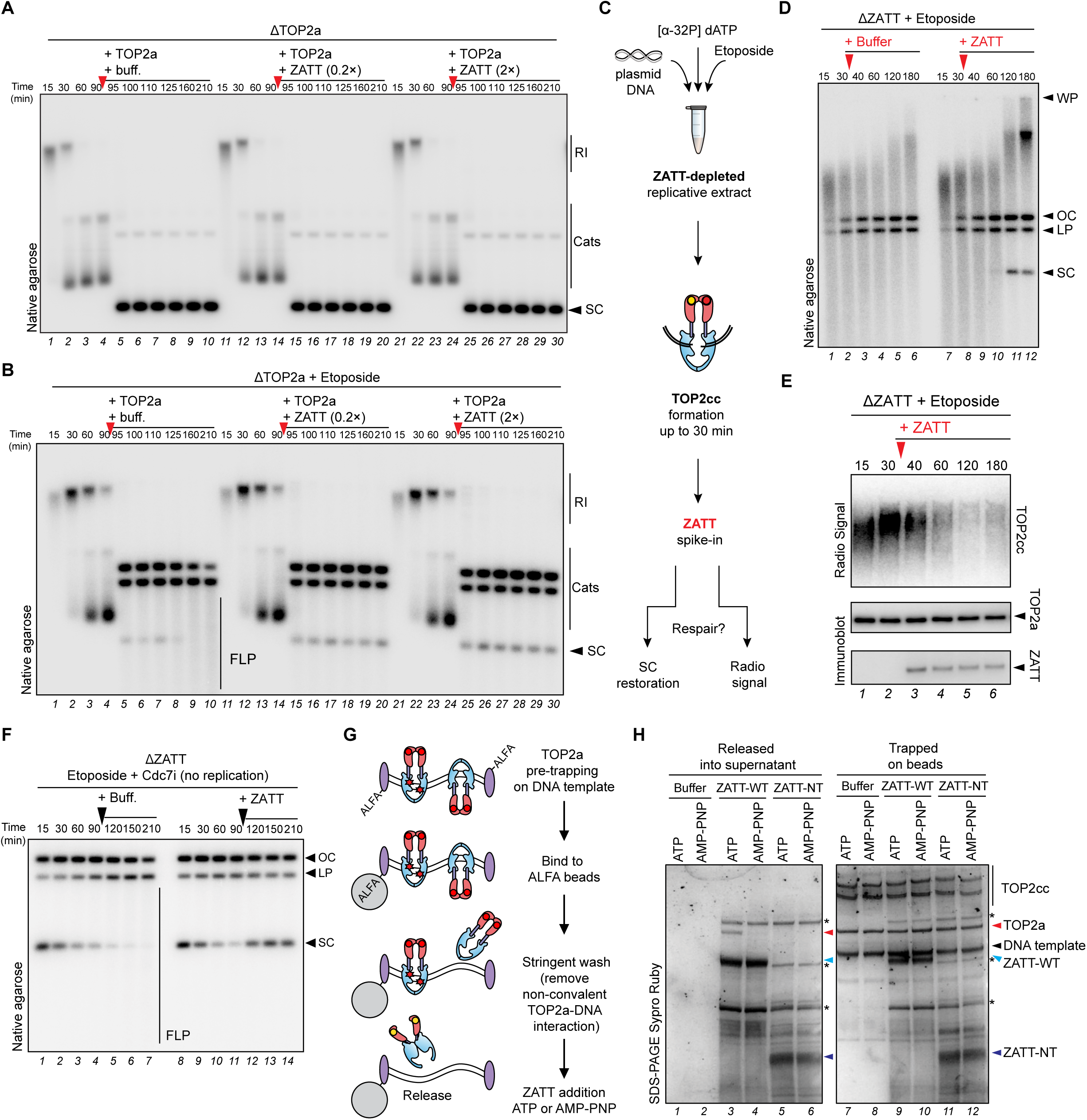
ZATT promotes TOP2cc resolution. **A)** pBS was replicated in TOP2a– depleted extracts. At 91 minutes, reactions were spiked in at indicated timepoints (red arrows) with TOP2a or TOP2a in combination with ZATT. Reactions were analyzed as in Fig.1C. **B**) Same as (A) but with etoposide (100 µM) supplemented to the reaction. **C**) Illustration of experimental design of (D), where TOP2cc was formed in ZATT-depleted extract and ZATT was spiked in after 30 min of reaction. **D**) pBS was replicated in the presence of etoposide in ZATT-depleted extracts. At 31 minutes, either buffer or recombinant ZATT was supplemented to the reaction (red arrow). Samples were analyzed as in Fig.1C. **E**) Samples from (D) were analyzed by drag-on blot and immunoblots with indicated antibodies. **F**) Pre-labelled pBS was incubated in ZATT-depleted extract supplemented with etoposide and Cdc7i to block DNA replication. Recombinant ZATT was spiked into the reaction at 91 min. Samples were analyzed as in Fig.1E. Note that ZATT addition restored the emergence of repaired product SC. **G**) Illustration of TOP2cc *in vitro* resolution assay. **H**) TOP2cc DNA-bead complexes were incubated with buffer, recombinant ZATT wildtype (WT), or recombinant ZATT-NT for 30 minutes in the presence of ATP or non-hydrolysable ATP analog AMP-PNP where indicated. The supernatant (released) or beads (trapped) were recovered and analyzed via SDS PAGE followed by SYPRO Ruby staining. Red arrow indicates TOP2a, for which the release into the supernatant is ATP dependent. Blue arrow indicates ZATT-WT and purple arrow, ZATT– NT. * denotes non-specific protein.

To clearly test that ZATT promotes TOP2cc repair, we performed a ZATT spike-in experiment, where TOP2cc was pre-formed in the absence of ZATT and repair was monitored upon ZATT addition (**Fig.5C**). Specifically, plasmids were first replicated in ZATT-depleted extracts with etoposide to generate a high number of TOP2css for 30 min (evidenced by the formations of FLPs), and recombinant ZATT or buffer was subsequently added at 31 minutes. While buffer addition maintained FLPs, ZATT supplementation led to progressive resolution of FLPs and generation of fully repaired SC products at later time points (**Fig.5D**, lanes 9-12). This was accompanied by a concomitant reduction in the drag-on blot signal (**Fig.5E**). Notably, ZATT addition after TOP2cc formation also restored SC products in the replication-independent assay (**Fig.5F**). These results indicate a direct role for ZATT in stimulating TOP2cc repair.

### ZATT promotes ATP-dependent resolution of TOP2ccs

Our findings imply that ZATT promotes TOP2cc resolution independently of TDP2 and canonical DSB repair pathways, pointing to a direct role in repair. To test whether ZATT can stimulate repair on its own, we reconstituted the reaction *in vitro* (**Fig.5G**). Recombinant TOP2a was trapped on linear DNA immobilized on ALFA-tag beads using etoposide, followed by stringent washes to remove non-covalently trapped TOP2a (Fig.S5B-C). The resulting TOP2a-DNA complexes were then incubated with recombinant ZATT to monitor resolution.

As shown in Figure 4D, recombinant ZATT released pre-trapped TOP2a from the beads– DNA substrate, whereas buffer alone or the truncated ZATT-NT did not (**Fig.5H**, compare lane 3 to 1 and 5). Although a significant fraction of TOP2a was released, a large amount remained bead-bound, possibly due to re-binding of TOP2a to the DNA substrate after release, resulting in a dynamic equilibrium between trapped and free states (**Fig.5H**, lanes 7-12). Adding competitor DNA increased release and reduced bead-bound TOP2a (Fig.S5D), supporting this model. Given that ZATT-mediated TOP2cc repair does not require DNA DSB repair, we speculated that the DSB in TOP2cc is sealed by TOP2 itself via resumption of its catalytic cycle, which is ATP hydrolysis-driven. To test this hypothesis, we tested whether ZATT-mediated release of TOP2a requires ATP hydrolysis. Addition of ATP to the reaction promoted TOP2a release, whereas the non-hydrolysable analog AMP-PNP failed to do so (**Fig.5H**, compare lanes 3 and 4). Together, these results suggest that ZATT facilitates TOP2cc resolution via an ATP-dependent release mechanism, supporting a model of TOP2cc resolution by promoting TOP2a resumption of its ATP hydrolysis-driven catalytic cycle.

### A conserved ZATT-TOP2 interface regulates TOP2cc resolution

To investigate how ZATT recognizes and promotes direct repair of TOP2ccs, we employed AlphaFold3 to assess potential interactions between TOP2a and ZATT (Abramson et al. 2024). Given that structure predictions with segments of proteins increases the likelihood to detect interactions (C. Y. Lee et al. 2024; Bret et al. 2024), we first modeled individual domains of human ZATT in complex with a human TOP2a monomer (Table S5). Zinc fingers (ZFs) were folded either individually or grouped within their respective clusters (i.e. ZF1-4, 5-6, 7-9, and 10-11). This analysis revealed that the coiled-coil domain of ZATT is predicted with high confidence to interact with TOP2a (Table S5). Notably, this interaction is compatible with both the dimerization of TOP2a and the formation of a TOP2cc (**Fig.6A-B**). A similar high-confidence interaction is also predicted between the ZATT coiled-coil region and TOP2b (Table S5). These interactions appear conserved and are likewise predicted with high confidence in *Xenopus laevis* (Table S5).

**Figure 6.**
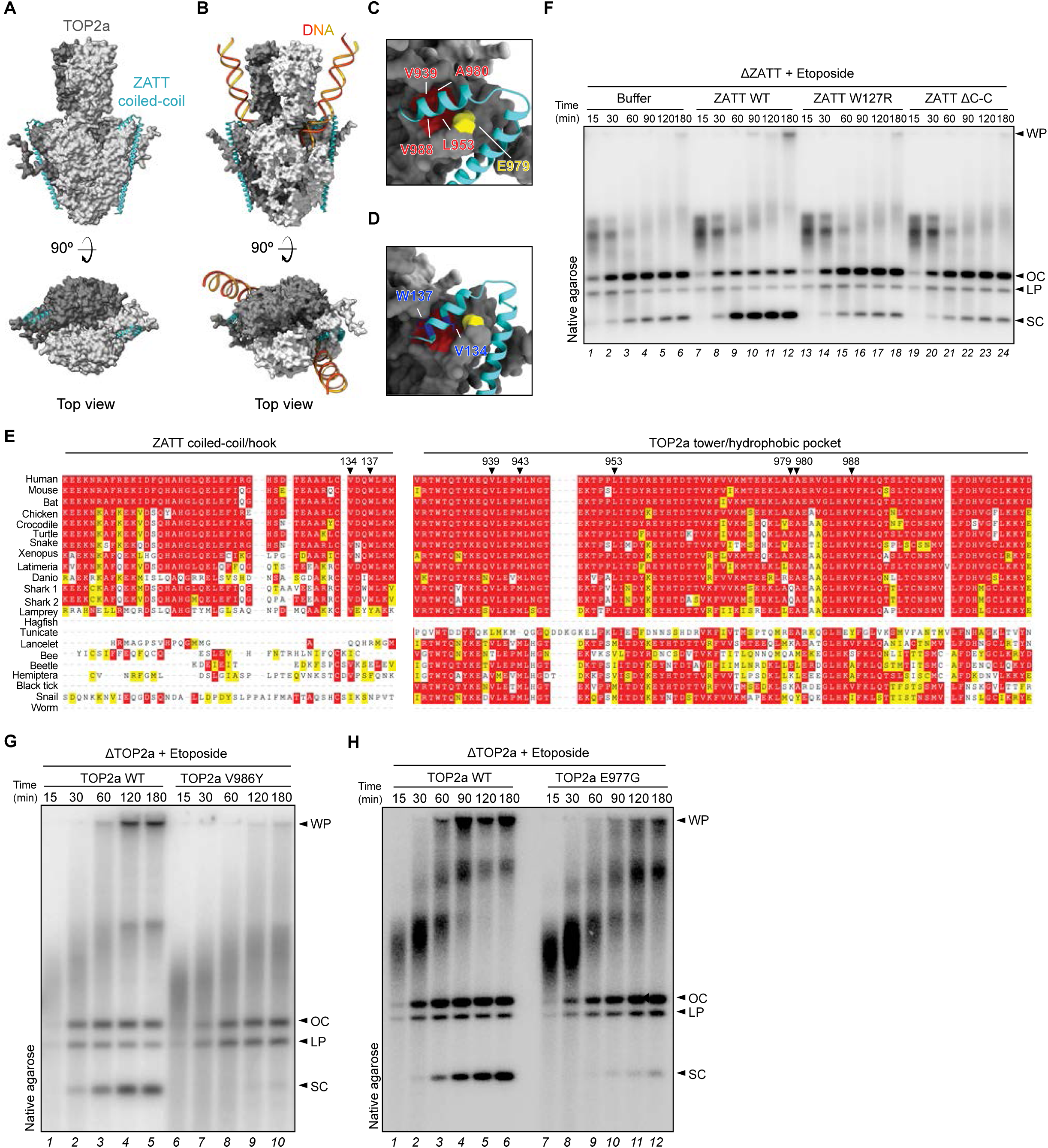
ZATT promotes TOP2cc resolution by binding to a hydrophobic pocket on TOP2a. **A**) AlphaFold3 predicted model of human TOP2a dimer (amino acids 40-1217) and ZATT coil-coiled domain (amino acids 60-140). **B**) AlphaFold3 predicted model of human TOP2acc and ZATT coil-coiled domain. **C**) Close up view of (A) highlighting the human TOP2a hydrophobic pocket residues (in red) and E979 (in yellow). **D**) Close up of (A) highlighting human ZATT W137 and V134 (in blue). **E**) Multiple sequence alignments of parts of the TOP2A and ZATT sequences from different species. Conserved residues are highlighted in red, conservative substitutions in yellow, and non-conservative substitutions in white. TOP2a residues located within the hydrophobic pocket and ZATT residues forming the tip of the hook-like motif are indicated with arrows. **F**) pBS was replicated in ZATT-depleted extracts in the presence of etoposide (100 µM). Where indicated, the reaction was supplemented with either *Xenopu*s ZATT WT, ZATT W127R, or ZATT ΔC-C. Samples were analyzed as in Fig.1C. **G**) pBS was replicated in TOP2a– depleted extracts in the presence of etoposide (100 µM). Reactions were supplemented with either *Xenopus* TOP2a WT or TOP2a V986Y. Samples were analyzed as in Fig.1E. **G**) pBS was replicated in TOP2a-depleted extracts in the presence of etoposide (100 µM). Reactions were supplemented with either *Xenopus* TOP2a WT or TOP2a E977G. Samples were analyzed as in Fig.1C.

Upon reanalyzing our tiling data and the guide RNAs targeting the ZATT coiled-coil domain, we identified two guides targeting the terminal end of the coiled-coil that conferred marked sensitivity to etoposide. These corresponded to predicted mutations at residues V134 and W137, with W137R ranking among the top 10 most sensitizing mutations in the screen (**Fig.4D**, Table S4). Structural predictions with AlphaFold3 suggest that these residues form the tip of a hook-like motif that inserts into a highly conserved hydrophobic pocket on the TOP2a tower domain, consisting of residues V939, M943, L953, A980, and V988 (**Fig.6C-E**), also conserved in TOP2b (composed of V960, M964, L974, A1001, and V1009) (Fig.S6A-D). ZATT W137 and V134 protrude into this pocket (**Fig.6E**), and the W137R mutation likely disrupts this interaction, providing a possible explanation for the observed etoposide hypersensitivity. The predicted hook wraps around an α-helix within the tower domain of TOP2a, with residue E979 of TOP2a forming key contacts with the ZATT coiled-coil (**Fig.6C**, Fig.S6E). Remarkably, neither a variant of ZATT lacking the coiled-coil (ΔC-C) or another carrying a W127R mutation (equivalent to human W137R) rescued the repair defect in ZATT-depleted extracts, confirming the essential role of this domain and residue (**Fig.6F**). To further test the ZATT– TOP2a interface, we engineered two TOP2a mutants. The first substituted a hydrophobic pocket residue (V988 in human corresponding to V986 in *Xenopus*) with tyrosine, and the second mutated E977 (equivalent to human E979) within the adjacent α-helix (Fig.S6F– G). Both mutants retained wild-type decatenation activity but failed to support repair after etoposide treatment, as indicated by loss of SC products (**Fig.6G-H**, S6H-I). These results underscore the critical role of the conserved hook-hydrophobic interface in ZATT– mediated TOP2cc resolution.

### The ZATT-TOP2 interface regulates TOP2cc repair in human cells

To validate the ZATT-TOP2a interface in human cells, we used a knockout and complementation system in U2OS Flp-In T-REx cells with doxycycline-inducible expression (Karvonen et al. 2008). ZATT-KO cells were complemented with eGFP-tagged ZATT WT, V134R, or W137R variants, all expressed at comparable levels and showing characteristic nuclear punctate localization (**Fig.7A**, Fig.S7A). ZATT-KO cells were hypersensitive to etoposide, which was rescued by ZATT WT but not by the V134R or W137R mutants (**Fig.7B-C**), supporting the functional importance of the conserved hook– hydrophobic interface.

**Figure 7.**
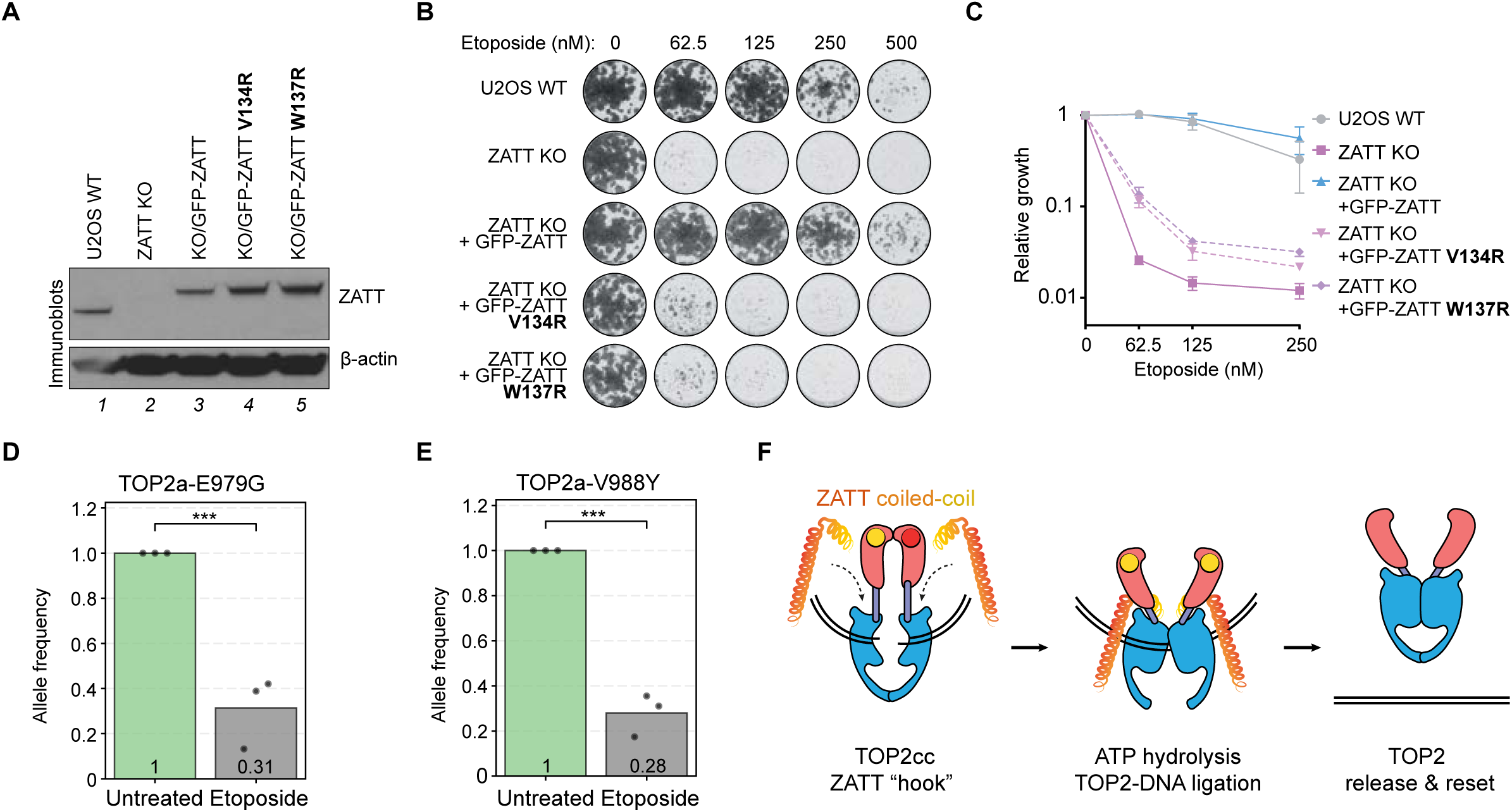
The ZATT-TOP2 interface regulates TOP2cc repair in human cells. **A**) Immunoblot analysis of whole cell extracts of the indicated cell lines. Samples were immunoblotted with the indicated antibodies. **B**) Representative images of SRB cell growth assay with indicated conditions and U2OS cell-lines. **C**) Quantification of SRB cell growth assay (mean±SD, n=3 biological replicates). **D**) CRISPR-Select analysis of TOP2a-E979G variant. CRISPR-Select cassettes were delivered to RPE P53 KO Flag– Cas9 cells, treated or untreated with etoposide for 12 days. Allele frequency was measured by NGS normalized to untreated value. Shown are means of 3 independent biological replicates. *** denotes p-value ≤ 0.001 by Student’s t-test. **E**) CRISPR-Select analysis of TOP2a-V988Y variant. Samples were analysed as in (D). **F**) Model illustrating ZATT-mediated TOP2cc resolution. The model highlights the ZATT “hook” binding to the TOP2 tower domain.

To functionally explore the role of TOP2a residues interacting with ZATT, we performed CRISPR-Select to test E979G and V988Y mutations (Niu et al. 2022). Cells were transfected with repair templates encoding either the mutation or a synonymous WT control, then cultured with or without etoposide for 10 days. Allelic frequencies were tracked via next-generation sequencing. Both TOP2a mutations conferred marked etoposide sensitivity (**Fig.7D-E**, Table S6), consistent with observations in *Xenopus* extract.

To determine whether this region essentiality is specific to ZATT-mediated repair, we assessed the effect of the TOP2a E979G mutation in a ZATT KO background (Fig.S7B). Notably, introduction of the E979G mutation into ZATT KO cells did not further sensitize these to etoposide, indicating genetic epistasis between TOP2a E979 and ZATT. Collectively, these findings confirm that the conserved hydrophobic pocket on TOP2a is essential for ZATT-mediated TOP2cc resolution and constitute a specific, potential target for therapeutic intervention.

## Discussion

Here we show that ZATT stimulates TOP2 release from its stalled cleavage complex (**Fig.7F**). Unlike other resolution pathways, ZATT-mediated TOP2cc resolution occurs without the induction of DSBs, making it an intrinsically safe mechanism. This provides a compelling explanation for the exquisite sensitivity of ZATT-deficient cells to TOP2 poisons. These insights not only deepen our understanding of genome maintenance but also open new avenues for chemotherapeutic strategies aimed at sensitizing cancer cells to TOP2 poisons.

### ZATT can function independently of TDP2

While ZATT was initially proposed to resolve TOP2ccs via TDP2 (Schellenberg et al. 2017), accumulating evidence highlights that ZATT also functions independently of TDP2 (Park et al. 2023; Schellenberg et al. 2017). Notably, ZATT KO cells exhibit greater sensitivity to etoposide than TDP2 KO cells, and ZATT-TDP2 double KO results in additive sensitivity (Park et al. 2023; Schellenberg et al. 2017). Accordingly, our findings in *Xenopus* egg extracts and *in vitro* reconstitution show how ZATT can resolve TOP2ccs in the absence of TDP2 (**Fig.2F-G**, **Fig.5H**), providing the mechanistic basis for these genetic observations.

Our *in vitro* experiments further show that ZATT-mediated TOP2 resolution requires ATP hydrolysis, suggesting that ZATT facilitates the resumption of TOP2’s catalytic cycle, thereby enabling TOP2 release from DNA. This represents the first evidence of a TOP2cc repair mechanism that operates independently of DSB exposure and repair, which we propose as the primary and inherently safe mechanism for TOP2cc resolution.

### A sequential model of TOP2cc repair

Why do cells maintain multiple orthogonal pathways to resolve TOP2ccs? We propose a sequential activation model in which ZATT acts as the primary safeguard against persistent TOP2ccs, promoting continuation of TOP2’s catalytic cycle and enabling its release from DNA without exposing the DSB (Fig.S7C, i). However, in situations where ZATT alone cannot resolve the lesion, either due to excessive DNA distortion or limited ZATT availability under high TOP2cc burden (e.g., etoposide treatment), secondary pathways become engaged. These include TDP2-mediated repair, first without proteolysis as previously proposed (Fig.S7C, ii) (Schellenberg et al. 2017). If this fails, proteolytic degradation of TOP2 followed by TDP2 or MRN-mediated nucleolytic incisions may be required, albeit at the cost of increased genome instability (Fig.S7C, iii) (Sciascia et al. 2020). In this study, we show how TOP2cc degradation via Trim-Away activates downstream DSB repair via MRN and NHEJ (**Fig.3**).

While TOP2a is abundantly expressed in human cells and in *Xenopus* egg extracts, ZATT appears to be present at significantly lower concentrations (Thul et al. 2017; Uhlen et al. 2017; Colding-Christensen et al. 2023). In cells, overexpression of ZATT alone confers resistance to etoposide (Schellenberg et al. 2017). If ZATT is so critical, why is it not more abundantly expressed? One possibility is that ZATT has roles beyond TOP2cc repair that must be carefully regulated (Karvonen et al. 2008; Y. Zhang et al. 2023). Moreover, endogenous TOP2ccs may rarely exceed the pool of ZATT molecules available for repair.

### ZATT binding to TOP2ccs

A recent study mapped the region of ZATT critical for its interaction with TOP2a (Park et al. 2023), identifying amino acids 1-168 as essential for co-immunoprecipitation in HEK293T cells. Our findings, supported by AlphaFold3 structural predictions, confirm a high-confidence interaction between ZATT’s coiled-coil domain (amino acids 60-139 in human ZATT) and the tower domain of TOP2a, an arrangement compatible with TOP2cc formation. CRISPR-based mutagenesis pinpoints key residues at the C-terminal end of ZATT’s coiled-coil domain whose disruption markedly increases etoposide sensitivity (**Fig.4B**). Functional validation through targeted mutations in *Xenopus* and human cells (**Fig.6**) confirms the importance of this anchoring interface.

AlphaFold3 further predicts that ZATT docks into a conserved hydrophobic pocket within the TOP2a TOWER domain, a feature also present in TOP2b (Fig.S6A-D), consistent with ZATT’s ability to resolve both TOP2 isoforms (Schellenberg et al. 2017). Importantly, a TOP2a mutant with alterations in this pocket retains decatenation activity but fails to be resolved from TOP2ccs, highlighting the pocket’s specific role in repair and its potential as a therapeutic target.

### The role of SUMOylation in TOP2cc resolution

Contrary to earlier models, our data show that ZATT’s SUMO ligase activity plays a minor role in TOP2cc repair. While earlier studies proposed that SUMOylation facilitates TDP2 recognition of TOP2ccs (Schellenberg et al. 2017), recent cellular evidence indicates that ZATT’s SUMO ligase function is dispensable for suppressing etoposide sensitivity (Park et al. 2023). Our tiling mutagenesis in cells and *Xenopus* egg extract experiments further support a limited contribution of ZATT’s SUMO ligase activity to TOP2cc repair (**Fig.4B**, **4F**).

Nonetheless, SUMO signaling appears to be a critical regulatory element in the repair process (**Fig.1G**). *Xenopus* egg extracts exhibit robust SUMO protease activity, as evidenced by the rapid loss of SUMO1/2/3 from plasmids upon SUMO inhibitor treatment (**Fig.2B**). SUMOylation enhances the recruitment of SUMO E3 ligases, including PIAS4 and ZATT, to chromatin (**Fig.2B**). Given that TOP2 is pre-SUMOylated on chromatin (Yoshida et al. 2016), this modification may be sufficient to enhance ZATT recruitment and initiate TOP2cc resolution, independent of ZATT intrinsic SUMO ligase activity. When ZATT-mediated resolution is insufficient, progressive poly-SUMOylation may act as a signal to trigger backup repair pathways, such as TDP2-dependent repair or STUbL– mediated proteasomal degradation (Schellenberg et al. 2017; Sun et al. 2020). Thus, while ZATT’s SUMO ligase function is not essential for its core repair activity, SUMOylation itself may still play a central role in coordinating the hierarchical activation of TOP2cc repair mechanisms.

### ZATT putative mechanism of action

Beyond its coiled-coil domain, our study reveals a critical role for ZATT’s zinc fingers (ZFs) in TOP2cc resolution. A clear functional gradient emerges: ZF1-4, located closest to the coiled-coil region, are most essential, followed by ZF5-6, and then ZF7-9 and ZF10-11. Notably, ZF1-4 are located near the tip of the “hook” that docks into the hydrophobic pocket of TOP2a, precisely where the G segment DNA exits the TOP2a dimer (**Fig.6B**). We propose that this docking positions ZF1-4 at the DNA exit site, enabling direct interaction with DNA and potentially pushing and/or stabilizing the DNA gate in a closed, ligation-competent state. ZF5-6 and the more distal ZFs may engage DNA regions further away from the DNA exit site, reinforcing the closed conformation and enhancing religation efficiency. While ZF7-11 appear dispensable for core repair activity in *Xenopus* extracts, tiling mutagenesis suggests they likely contribute under conditions where religation is challenging (**Fig.4B**).

A key mechanistic question is how ZATT promotes religation in the presence of etoposide. Etoposide is known to transiently dissociate from DNA (Burden et al. 1996; Wu et al. 2011; Kuriappan, Osheroff, and De Vivo 2019). By stabilizing a ligation-ready intermediate, ZATT may allow TOP2 to complete religation during these brief windows of etoposide disengagement, thereby resolving the TOP2cc.

### Therapeutic implications

Although widely used across multiple cancer types, etoposide treatment is associated with severe side effects, most notably bone marrow toxicity, which increases the risk of infection and anemia. These toxicities have significantly limited the dosage that can be safely administered, likely contributing to both the high incidence of treatment resistance and the emergence of secondary leukemias (Ezoe 2012).

Our findings suggest that targeting the hydrophobic pocket of TOP2, can selectively impair TOP2cc resolution via ZATT without inhibiting TOP2 activity. This strategy may enhance etoposide efficacy while allowing for reduced dosing, potentially mitigating its toxic side effects. Looking ahead, we envision novel antibody-drug conjugate (ADC) strategies that co-deliver etoposide with a small molecule inhibitor targeting the TOP2 pocket or disrupting ZATT’s coiled-coil interaction. Such dual-targeting ADCs could synergistically block TOP2cc repair and amplify cytotoxicity, offering a potent and selective therapeutic cocktail, a “death sentence” for cancer cells.

## Materials and methods

### Methods

#### *Xenopus* egg extracts and DNA replication reactions

*Xenopus* egg extracts were prepared as described previously (Lebofsky, Takahashi, and Walter 2009). All experiments involving animals were approved by the Danish Animal Experiments Inspectorate and conform to relevant regulatory standards and European guidelines.

For plasmid DNA replication, pBluescript II SK(+) (pBS) was licensed in high-speed supernatant (HSS) at a final concentration of 7.5 ng/µL for 30 minutes at room temperature (RT). Replication was initiated by adding two volumes of nucleoplasmic egg extract (NPE). Etoposide was supplemented at 100 µM (F.C.) to replication reactions and at 200 µM (F.C.) to non-replication reactions. For non-replication reactions, NPE was incubated with 150 µM of Cdc7i (Sigma-Aldrich) for 10 minutes at RT prior to addition to the licensing reaction.

The Ubiquitin activating enzyme inhibitor (Ubi) MLN-7243 (MedChem Express), SUMOylation activating enzyme inhibitor (SUMOi) ML792 (MedKoo Biosciences), or proteasome inhibitor MG262 (Boston Biochem) were supplemented to NPE at a final concentration of 200 µM, 10 minutes before initiating the reaction.

Etoposide (Sigma-Aldrich) or ICRF-193 (Enzo Life Sciences) were supplemented in NPE at the indicated final concentrations, 10 minutes before initiating the reaction.

To visualize DNA replication intermediates, replication reactions were supplemented with [α-^32^P]dATP (Perkin Elmer). For each timepoint, 1 µL of the reaction mixture was added to 5 µL of stop buffer (5% SDS, 80 mM Tris pH 8.0, 0.13% phosphoric acid, 10% Ficoll), followed by addition of 1 µL of Proteinase K (20 mg/mL) (Roche). The samples were incubated at 37°C for 1 hour and subsequently separated by using 0.9% native agarose gel electrophoresis, and visualization using a phosphor imager. The radioactive signal was quantified using ImageJ (NIH, USA).

To visualize reactions in the absence of DNA replication, plasmids were first radiolabeled *in vitro* via nick translation synthesis. Briefly, pBS was nicked with Nb.BsrDI (NEB). The DNA was subsequently radiolabeled with [α-^32^P]dATP via nick translation synthesis by DNA pol I (NEB) for 20 minutes at 16 °C.

### Antibodies and Immuno-depletions

Antibodies against ORC2 (Fang and Newport 1993), BRCA2 (Ambjørn et al. 2021) were described previously. Antibody against 6×HIS was purchased (Fisher Scientific).

ZATT, TOP2a, TDP2, XRCC4, MRE11, Polθ were raised and affinity purified by New England Peptide by immunizing rabbits with Ac-CDEDANLNEAIKRSMTEM-OH (ZATT), Ac-CAQAGRQKKPVTYLEDSDDDF-OH (TOP2a), Ac-CKNTPDPDDLFSDI-OH (XRCC4), Ac-CDDEEDFDPFKKSGPSRRGRR-OH (MRE11). RAD51 antibody was raised and affinity purified by Proteogenix by immunizing rabbits with CAEAMFAINADGVGDAKD.

To immunodeplete ZATT, TOP2a, TDP2, XRCC4, MRE11, Polθ, and BRCA2 from *Xenopus* egg extracts, corresponding antibodies (1mg/mL) were first coupled to Protein A Dynabeads™ (Thermo Fisher Scientific) in 1:3 ratio overnight at 4°C. Beads then were washed with PBS buffer (with 0.02% NaN_3_) and ELB buffer (25 mM MgCl_2_, 0.5 M KCl, 100 mM HEPES, pH7.7). Beads were washed and dried on magnetic stand. Dried beads were incubated with HSS or NPE at 3:1 ratio for 25 min on a rotating wheel to deplete target proteins. Immuno-depletion was performed two rounds, NPE or HSS was then separated from beads on magnetic stands. Immuno-depleted NPE or HSS was subsequently used in replication or non-replication reactions. For depletion and addback reactions, recombinant ZATT was added to 60 nM (F.C.) in replication reactions and 120 nM (F.C.) in replication-independent reactions. TOP2a was added to 500 nM (F.C.) to rescue TOP2a-depletions in both replication and replication-intendent reactions.

### Cloning, protein expression, and purification

#### Generation of Xenopus TOP2a expression constructs

*Xenopus* TOP2a WT expression plasmids were kindly provided by Prof. Yoshiaki Azuma (University of Kansas, Lawrence). *Xenopus* TOP2a mutants were generated using Q5 Site-Directed Mutagenesis Kit (NEB) following manufacturer’s protocol. Primers used to generate TOP2a mutants are summarized in Table S7.

#### Xenopus TOP2a expression and purification

Recombinant TOP2a proteins were purified from budding yeast (JEL-1), which was transformed with target TOP2a expression construct using Frozen-EZ Yeast Transformation II Kit (Zymo Research) following manufacturer’s protocol. Transformed yeast grew on SC-URA agar plates for three days and 10 colonies were picked and inoculate in 20 mL SC-URA (FORMEDIUM) media with 2% glucose (FORMEDIUM) for 48 hours. Yeast culture was then transferred into 300 mL SC-URA media with 2% raffinose (FORMEDIUM) and grew for 24 hours. Culture was diluted to OD_600_=0.3 into YEP (FORMEDIUM) media with 2% raffinose. The yeast grew at 30°C with shaking until OD_600_=0.8. The yeast culture was supplemented with final 2% galactose (FORMEDIUM) and temperature was decreased to 20°C to induce expression of recombinant TOP2a. After 24 hours, yeast cells were collected by centrifugation and snap-frozen into liquid nitrogen with PBS buffer.

To purify FLAG-tagged recombinant TOP2a, frozen yeast cells were ground using freezer mill and thawed on ice, followed by high-speed centrifugation at 25000×g for 30 minutes at 4°C. After centrifugation, the supernatant was mixed with 1 mL/3L culture of ANTI– FLAG® M2 Affinity Gel (Sigma-aldrich) and incubated at 4°C for 90 minutes. ANTI-FLAG® M2 Affinity Gel was then transferred into a 25 mL gravity column (Bio-rad) and washed four times with 25 mL of high salt buffer (50 mM Tris-HCl, 500 mM NaCl, 10% Glycerol, 1 mM EDTA, 0.1 mM DTT and Roche EDTA-Free Protease Inhibitor) and twice with 25 mL of low salt buffer (50 mM Tris-HCl, 200 mM NaCl, 10% Glycerol and 1 mM EDTA). FLAG– tagged recombinant TOP2a was eluted from ANTI-FLAG® M2 Affinity Gel using 0.33 mg/mL 3×FLAG peptide (ApexBio) in low salt buffer. Eluted recombinant TOP2a was concentrated using protein spin-column concentrator (Sartorious) and were snap-frozen in liquid nitrogen and stored at –80°C.

#### Generation of Xenopus ZATT (ZNF451) expression constructs

*Xenopus* ZNF451 (ZATT) full length cDNA (Horizon) was cloned into expression construct p0197 (NNF-CPR). *Xenopus* ZATT mutants were generated using Q5 Site-Directed Mutagenesis Kit (NEB) following manufacturer’s protocol. Primers used to generate ZNF451 mutants are summarized in Table S7.

#### Xenopus ZATT (ZNF451) expression and purification

Recombinant ZATT proteins were purified from HEK293-6E suspension cells. HEK293– 6E cells were maintained at 1 million/mL in FreeStyle F17 expression medium (Gibco) supplemented with 4 mM L-glutamine (Thermo Scientific), 1% FBS (Gibco), 0.1% Poloxamer 188 (Gibco) and 50 μg/ml G418 (InvivoGen). To transfect 1 million cells, 1 µg of ZATT expression construct DNA was mixed with 2 µg of PEI-MAX (Polyscience) in Opti-MEM medium (Gibco) and incubate at room temperature for 20 minutes. DNA/PEI– MAX mixture was then added to cells. 24 hours post transfection, valproic acid (Sigma Aldrich) was added to cell culture at a final concentration of 3.75 mM to enhance expression of target protein. Cells were harvested 48-72 hours post transfection by centrifugation at 1000×g for 10 minutes, pellet was washed with PBS.

To purify Strep-tagged recombinant *Xenopus* ZNF451 (ZATT), cell pellet from 1 L culture was resuspended in 30 mL resuspension buffer (350 mM NaCl, 50 mM Tris pH=8, 0.05% NP-40, 100 µM ZnSO_4_, Roche EDTA-Free Protease Inhibitor and 1 mM DTT) and lysed via sonication (40% amplitude with 1 second on and 1 second off for 3 minutes in total) with presence of 4 µL Benzonase (Merck-Millipore). The lysate was then spun down at 22000×g at 4 °C for 30 minutes. Recovered supernatant was transferred to a new tube and mixed with 1.5 mL of Strep-Tactin® 4Flow® resin (IBA Lifesciences) and incubated on a tube roller for 2 hours at 4°C. After incubation, the mixture was transferred into a gravity column (Bio-Rad), and washed twice with 25 mL resuspension buffer, twice with 25 mL washing buffer (600 mM NaCl, 50 mM Tris pH=8, 0.05% NP-40, 100 µM ZnSO_4_, Roche EDTA-Free Protease Inhibitor and 1 mM DTT), twice with 25 mL ATP buffer (600 mM NaCl, 5 mM ATP, 5 mM MgCl2, 50 mM Tris pH=8, 0.05% NP-40, 100 µM ZnSO4, Roche EDTA-Free Protease Inhibitor and 1 mM DTT) and once with 25 mL resuspension buffer. Strep-tagged ZATT was eluted with 3.5 mL (7 fractions of 500 µL) elution buffer (100 mM Tris pH=8, 350 mM NaCl, 1 mM DTT, and 5 mM Desthiobiotin). Eluted proteins were concentrated with using protein spin-column concentrator (Sartorious) and were snap-frozen in liquid nitrogen and stored at –80°C.

### Cell-free Trim-Away assay in *Xenopus* egg extracts

TRIM21 expression construct was a kind gift from Prof. Johannes Walter (Harvard Medical School). TRIM21 protein was expressed and purified as previously described (Mevissen, Prasad, and Walter 2023).

Cell-free Trim-Away assay was performed largely as previously described to apply to reactions in *Xenopus* egg extracts (Mevissen, Prasad, and Walter 2023). Specifically, a 10× TRIM21-antibody mixture containing 10 μM TRIM21 and 5 μM of TOP2a antibodies was incubated for 10 minutes at room temperature before adding to the reactions. At indicated timepoint, 10× TRIM21-antibody mixture was added 1:10 to the reactions to trigger TOP2cc degradation.

### DNA Relaxation Assay

DNA relaxation assay was performed in TTB-ATP buffer (50 mM Tris-HCl pH=8, 25 mM NaCl, 10 mM MgCl_2_, 2 mM ATP, 1 mM DTT) supplemented with plasmid pBluescript II SK(+) (pBS), recombinant ZATT and TOP2a at indicated concentration. 15 µL reactions were stopped at indicated timepoints by adding 15 µL 2× Gel Loading Dye Purple (NEB), followed by the addition of 1 µL of Proteinase K (20 mg/mL) (Roche). The samples were incubated at 37°C for 1 hour and separated using 1% native agarose gel electrophoresis. The gel was stained using SYBR Gold (Invitrogen) following manufacture’s protocol and imaged using Amersham ImageQuant 800 (Cytiva).

### Drag-on blotting

A DNA-protein crosslink drag-on blotting assay was developed to visualize and quantify the correlation between radiolabeled DNA crosslinked to proteins. To perform Drag-on blotting, samples containing [α-32P]dATP incorporated DNA-protein crosslink were denatured in 2×sample buffer and boiled at 95°C for 5 minutes. Samples were then separated by SDS-PAGE on gradient Criterion™ Protein Gel (BIO-RAD) and transferred onto a PVDF membrane via standard western blotting procedure. The PVDF membrane was briefly dried with Wattman paper and exposed in an exposure cassette. The signal of [α-32P]dATP incorporated DNA was visualized using a phosphor-imager and quantified by ImageJ. The same PVDF membrane bound was subsequently blotted with a TOP2a antibody to visualize TOP2a protein migration.

To quantify the relative TOP2cc signal, the radiolabeled signal detected on the PVDF membrane was normalized to the total [α-32P]dATP incorporation on the plasmid quantified on a native agarose gel.

### In-vitro TOP2cc resolution assay

To test TOP2cc resolution in-vitro, purified recombinant TOP2a was pre-trapped on a fluorinated double-stranded DNA, which was generated as previously described (Miller et al. 2019). Briefly, the DNA substrate was generated by PCR amplification. The PCR product was purified by anion exchange chromatography on a RESOURCE Q 1 mL column (Cytiva) equilibrated in 50 mM Tris-HCl pH=8, 5 mM beta-mercaptoethanol. The amplified PCR products were eluted with a linear gradient from 100 mM to 1 M NaCl at 2 mL/min. Fractions containing the DNA were analyzed by agarose gel electrophoresis, pooled and DNA was precipitated by isopropanol addition to 70% v/v for 30 min at 14000 g at RT. The pellet was washed with 70% ethanol for 1h at 4 degrees by centrifugation at 14000 g and resuspended in 10 mM Tris-HCl pH=8, 1 mM EDTA for 30 min at 23 degrees in benchtop shaker at 550 rpm. To carry out the conjugation reaction, the resuspended DNA was supplemented with an 8-fold molar excess of ALFA-tagged methyltransferase (M.HpaII) and 0.15 µM S-Adenosyl methionine (SAM). After incubating for 30 min at 37 °C on shaker at 450 rpm, 1.4 uL SAM per 300 µL reaction were spiked in before leaving overnight at 30 °C. The conjugated DNA product was further purified by anion exchange chromatography with the same column and gradient as the PCR product. The fractions containing the conjugated DNA were pooled, dialyzed twice at 4 °C against 2L ELB buffer using 500 µL slide-a-lyzers with 10 kDa MWCO (Thermo Fisher) and concentrated with a 30 kDa MWCO 0.5 mL centricon (Millipore) at 13500 g, fixed rotor, 4 °C. To confirm the purity and conjugation extent, the conjugated DNA was analyzed by agarose gel electrophoresis in the presence or absence of proteinase K treatment.

To generate TOP2accs, recombinant TOP2a was mixed at a final concentration 0.32 µM with 0.32 µM of the double strand template DNA and etoposide at a final concentration 500 µM in TOP2-trapping buffer (50 mM Tris-HCl pH=8, 25 mM NaCl, 10 mM MgCl_2_ and 1 mM DTT) supplemented with either 2 mM ATP or AMP-PNP. The TOP2a pre-trapping reaction was incubated at room temperature for 30 minutes. After trapping, TOP2cc was bound to 40 µL ALFA Selector PE (Nanotag Biotechnologies) with etoposide at a final concentration 500 µM in TOP2-trapping buffer (50 mM Tris-HCl pH=8, 25 mM NaCl, 10 mM MgCl2 and 1 mM DTT) supplemented with either 2 mM ATP or AMP-PNP. After binding, ALFA selector beads were washed twice in TOP2 trapping buffer containing 250 mM NaCl, to remove unbound Top2a. Beads were then dried on a magnetic stand and added into TOP2cc release solution containing 0.32 µM recombinant ZATT in TOP2– trapping buffer (50 mM Tris-HCl pH=8, 25 mM NaCl, 10 mM MgCl_2_ and 1 mM DTT) supplemented with either 2 mM ATP or AMP-PNP. To reduce TOP2a re-engagement on DNA, external bait DNA pBS plasmid (at indicated molar ratio) was added into release reactions. The release reactions were gently mixed on a rotating wheel at room temperature for 30 minutes. The reactions were spun down at 500×g for 1 minute and flow-through supernatant was recovered into a new tube with 2×sample buffer. The remaining ALFA Selector beads were resuspended in 2×sample buffer. Protein samples from flow-through (released fraction) and beads (un-released fraction) were boiled at 95°C for 5 minutes and subsequently used in SDS-PAGE analysis. The SDS-PAGE gel was analyzed by Sypro Ruby Protein Blot Stain kit (Thermo Fisher) following manufacturer’s protocol.

### In vitro SUMOylation reaction

To monitor ZATT auto-SUMOylation activity, recombinant wild-type ZATT and ZATT mutant variants were incubated at indicated concentration (i.e. 0.5 µM, 1 µM and 2 µM) with 100 ng human SAE1 and UBA2 (Boston Biotech), 300 ng UBC9 (Boston Biotech) and 3 µg SUMO2 (Boston Biotech) per 10 µl reaction in 1×Buffer S (50 mM HEPES pH=7.5, 100 mM NaCl, 10 mM MgCl_2_, 100 nM DTT, 2 mM ATP and 0.2 mg/mL BSA). Reaction was performed at room temperature for 60 minutes and stopped by adding 2×sample buffer. *In vitro* SUMOylation was detected using anti-6×His (Fisher Scientific) or anti-SUMO2/3 antibody (Abcam) by immunoblotting.

### Plasmid Pull-down Mass Spectrometry (PP-MS)

Plasmid Pull-down Mass Spectrometry was conducted previously described (Larsen et al. 2019). Briefly, plasmid DNA was replicated in egg extracts at 5 ng/µL (final concentration). At the indicated time points, 8 µL of the reaction were withdrawn and plasmids and associated proteins were recovered by plasmid pull down using LacI coated beads. After 30 min incubation at 4°C, samples were washed twice in 10 mM HEPES pH 7.7, 50 mM KCl, 2.5 mM MgCl2, 0.03% Tween 20, and once in 10 mM HEPES pH 7.7, 50 mM KCl, 2.5 mM MgCl2. Samples were washed one additional time in 50 µL of 10 mM HEPES pH 7.7, 50 mM KCl, 2.5 mM MgCl2 and transferred to a new tube to remove residual detergent. Beads were dried out and resuspended in 50 µL denaturation buffer (8 M Urea, 100 mM Tris pH 8.0). Cysteines were reduced (1 mM DTT, 15 minutes at RT) and alkylated (5 mM iodoacetamide, 45 min at RT). Proteins were digested and eluted from beads with 1.5 µg LysC (Sigma) for 2.5 hr at RT. Eluted samples were transferred to a new tube and diluted 1:4 with ABC (50 mM ammonium bicarbonate). Subsequently, 2.5 µg trypsin was added and incubated for 16 hours at 30°C, after which NaCl was added to a final concentration of 400 mM and samples were cleaned up using C18 StageTips.

### Total proteome mass spectrometry (TP-MS)

Xenopus nucleoplasmic egg extracts (NPE) were either mock– or TDP2-depleted (see above). Subsequently, digestion of extracts was performed using modified sequencing grade Trypsin (1:200 w/w; Sigma) for 30 min at room temperature, followed by 10-fold dilution with 50 mM ammonium bicarbonate. Samples were reduced and alkylated for 30 min at room temperature by addition of Tris-(2-Carboxyethyl)phosphine (TCEP) and chloroacetamide (CAA) to final concentrations of 10 mM, after which additional Trypsin (1:200 w/w) was added and digestion was allowed to continue overnight. Digests were clarified by centrifugation through 0.45 µm spin filters prior to C18 StageTip cleanup.

### Immunoprecipitation mass spectrometry (IP-MS)

Ether control IgG, anti-TOP2a or anti-ZATT antibody was linked to Protein A Sepharose resin overnight at 4 °C at a beads:antibody ratio of 1:3. Beads were then washed in cold PBS, cold ELB buffer, ELB-high salt (0.5 M NaCl), and finally kept in ELB buffer until immunoprecipitation. After 60 minutes, one volume of Etoposide-treated replicating egg extract reaction was then added to 0.4 volumes of antibody-bound beads in a total of 5 volumes of IP buffer (ELB buffer supplemented with 0.3 % NP-40, RNase A and Benzonase) and incubated at 4 °C for 2 h with end-over-end rotation. After the incubation, resin was washed 3 times with 0.5 mL of ELB-High Salt + 0.25% NP-40, once with 0.5 mL of ELB-High Salt, and transferred to LoBind tubes. Subsequently, resin was washed 3 times with 0.5 mL of ELB-High Salt, and beads were dried using ultra-fine tips. 50 µL of ice-cold 50 mM ammonium bicarbonate containing Trypsin (10 ng/mL) was added, and resin was pre-incubated for 30 min at 4 °C while shaking, prior to overnight shaking at 30 °C. Digests were clarified by centrifugation through 0.45 µm spin filters, priortp reduction and alkylation for 30 min at room temperature by addition of TCEP and CAA to final concentrations of 10 mM, and C18 StageTip cleanup.

### Peptide clean-up for mass spectrometry

C18 StageTips were prepared in-house, by layering four plugs of C18 material (Sigma– Aldrich, Empore SPE Disks, C18, 47 mm) per StageTip. TP-MS and PP-MS samples were processed at high pH, whereas IP-MS samples were processed at low pH. For high pH (TP-MS and PP-MS), activation of StageTips was performed with 100 μL 100% methanol, followed by equilibration using 100 μL 80% acetonitrile (ACN) in 50 mM ammonium hydroxide, and two washes with 100 μL 50 mM ammonium hydroxide. Samples were basified to pH >11 by addition of ammonium hydroxide to a final concentration of 200 mM, centrifuged for 5 min at 14,000*g*, after which supernatants were loaded on StageTips. Subsequently, StageTips were washed twice using 100 μL 50 mM ammonium hydroxide. For PP-MS samples, the peptides were eluted using 80 µL 25% ACN in 50 mM ammonium hydroxide. For TP-MS samples, on-StageTip fractionation was performed by sequential elution using 40 µL of 2%, 4%, 6%, 9%, 12%, 16%, and 20% ACN in 50 mM ammonium hydroxide. For low pH (IP-MS), processing was done as above but 0.1% formic acid was used instead of 50 mM ammonium hydroxide, and samples were acidified prior to centrifugation and loading by addition of trifluoroacetic acid (TFA) to a final concentration of 1%. Elution was done with 80 µL 30% ACN in 0.1% formic acid. All samples and fractions were dried to completion using a SpeedVac at 60 °C. Dried peptides were dissolved in 10 μL (PP-MS and IP-MS) or 20 µL (TP-MS) 0.1% formic acid. For IP-MS samples, half of the peptides were subjected to additional cleanup using SCX StageTips, which were prepared in-house by layering four plugs of SCX material (Sigma– Aldrich, Empore SPE Disks, Cation, 47 mm) per StageTip. SCX StageTips were activated using 50 µL of 100% ACN, after which peptides were acidified by addition of TFA to a final concentration of 1% and loaded. SCX StageTips were washed twice using 100 µL of 40% ACN in 0.15% TFA, and eluted using 100 µL of 80% ACN in 200 mM ammonium hydroxide. SCX elutions were dried to completion using a SpeedVac at 60 °C, and dissolved in 10 µL 0.1% formic acid. All samples were stored at −20 °C until analysis using mass spectrometry (MS).

### MS data acquisition

TP-MS and PP-MS samples were analyzed on an EASY-nLC™ 1200 system (Thermo) coupled to an Orbitrap™ Exploris™ 480 mass spectrometer (Thermo), whereas IP-MS samples were analyzed on a Vanquish™ Neo UHPLC system (Thermo) coupled to an Orbitrap™ Astral™ mass spectrometer (Thermo). For TP-MS, 35% to 62.5% of each fraction was analyzed to achieve equal (∼1 µg) peptide loading. For PP-MS, 50% of each sample was analyzed. For IP-MS, 5% of the C18-cleaned samples were analyzed, whereas 15% of the C18-SCX-cleaned samples were analyzed. All samples were analyzed on 20 cm long analytical columns, with an internal diameter of 75 μm, and packed in-house using ReproSil-Pur 120 C18-AQ 1.9 µm beads (Dr. Maisch). Elution of peptides from the column was achieved using a gradient ranging from buffer A (0.1% formic acid) to buffer B (80% acetonitrile in 0.1% formic acid), at a flow rate of 250 nL/min.

Gradient length was 80 min (PP-MS and TP-MS), 40 min (IP-MS C18-SCX), or 30 min (IP-MS C18) per sample, including ramp-up and wash-out, with a primary analytical gradient of 60 min, 30 min, or 22 min, respectively. The primary analytical gradient increased in % of buffer B as follows: 5-37 (PP-MS), 3-24 (TP-MS F1), 4-26 (TP-MS F2), 4-28 (TP-MS F3), 4-30 (TP-MS F4), 4-32 (TP-MS F5), 5-34 (TP-MS F6), 5-36 (TP-MS F7), 5-40 (IP-MS C18-SCX), and 5-38 (IP-MS C18). The column was heated to 40 °C using a column oven (Sonation), and ionization was achieved using a NanoSpray Flex™ NG ion source (Thermo). Spray voltage set at 2 kV, ion transfer tube temperature to 275°C, and RF funnel level to 40% (PP-MS and TP-MS) or 50% (IP-MS). All full precursor (MS1) scans were acquired using the Orbitrap™ mass analyzer, with full scan range set to 300-1,300 m/z, MS1 resolution to 120,000 (PP-MS and TP-MS) or 180,000 (IP-MS), MS1 AGC target to 2,000,000 charges (PP-MS and TP-MS) or 5,000,000 charges (IP– MS), and MS1 maximum injection time to “Auto” (PP-MS and TP-MS) or 150 ms (IP-MS). Precursors were analyzed in data-dependent acquisition (DDA) mode, with charges 2-6 selected for fragmentation using an isolation width of 1.3 m/z and fragmented using higher-energy collision disassociation (HCD) with normalized collision energy of 25. Monoisotopic Precursor Selection (MIPS) was enabled in “Peptide” mode.

Repeated sequencing of precursors was minimized by setting dynamic exclusion duration to 60 s (PP-MS and TP-MS), 15 s (IP-MS C18-SCX), or 10 s (IP-MS C18), with an exclusion mass tolerance of 15 ppm (PP-MS and TP-MS) or 10 ppm (IP-MS) and exclusion of isotopes. MS2 scans were acquired using the Orbitrap mass analyzer on the Exploris MS (PP-MS and TP-MS), and the Astral mass analyzer on the Astral MS (IP– MS). MS2 fragment scanning method was set to first mass 100 m/z (PP-MS and TP-MS) or scan range 100-1,500 m/z (IP-MS), and MS2 AGC target to 200,000 charges (PP-MS and TP-MS) or 10,000 charges (IP-MS). On the Exploris (PP-MS and TP-MS), MS2 scans were recorded at a resolution of 45,000 (PP-MS) or 15,000 (TP-MS), intensity threshold was set to 230,000 (PP-MS) or 430,000 (TP-MS) charges per second, MS2 maximum injection time was set to “Auto” (PP-MS and TP-MS), and 9 (PP-MS) or 18 (TP-MS) data– dependent scans were performed per cycle. On the Astral (IP-MS), MS2 scans were recorded using the Astral analyzer (resolution 80,000), and were acquired at two priorities, first and second, with MS2 intensity threshold set to 100,000 and 25,000 charges per second, and MS2 maximum injection time to 5 and 20 ms, respectively. Cycle time was fixed at 0.5 s.

### MS data analysis

MS RAW data were analyzed using MaxQuant software v1.5.3.30 for the PP-MS and TP– MS datasets, and v.2.5.2.0 for the IP-MS dataset. The three datasets (PP-MS, TP-MS, and IP-MS) were computationally processed separately. Default MaxQuant settings were used, with exceptions specified below. For generation of theoretical spectral libraries, the Xenopus laevis FASTA database was downloaded from UniProt on the 3^rd^ of October 2022 and contains 67,462 entries. In-silico digestion of proteins to generate theoretical peptides was performed with trypsin, allowing up to 2 (PP-MS; default) or 3 (IP-MS) or 4 (TP-MS) missed cleavages. Carbamidomethylation of cysteine was defined as a fixed modification. Allowed variable modifications were oxidation of methionine (default), protein N-terminal acetylation (default), peptide N-terminal conversion of glutamine to pyroglutamate, deamidation of asparagine and glutamine (TP-MS and IP-MS), deamidation of asparagine (PP-MS), dioxidation of tryptophan (PP-MS), replacement of three protons by iron on glutamate and aspartate (PP-MS), and tryptophan oxidation to kynurenine (PP-MS). For all searches, the maximum variable modifications per peptide was set to 3. First search mass tolerance was set to 10 ppm (PP-MS and TP-MS) or 20 ppm (IP-MS; default), with main search mass tolerance set to 4.5 ppm (default). Matching between runs was enabled, with an alignment window of 20 (PP-MS and TP-MS) or 10 min (IP-MS) and a match time window of 1 (PP-MS and TP-MS) or 0.2 min (IP-MS). Label– free quantification (LFQ) was enabled, with “Fast LFQ” disabled. iBAQ quantification was enabled. Stringent MaxQuant 1% FDR data filtering at the PSM– and protein-levels was applied (default).

### MS data quantification and statistical analysis

All quantitative experiments were performed using at least four replicates to ensure sufficient statistical power. Statistical handling of the data was primarily performed using the freely available Perseus software (Tyanova et al. 2016) and includes scatter plot analysis and Pearson correlation (calculated using linear regression), volcano plot and heatmap analysis. For quantification purposes, all abundance values were Log_2_– transformed and filtered for presence in all four replicates in at least one experimental condition. Missing values were imputed below the global experimental detection limit at a downshift of 1.8 and a randomized width of 0.3 (in Log_2_ space; Perseus default). For the data presented in volcano plots, statistical significance of differences was tested using two-tailed Student’s t-testing, with permutation-based FDR control at 5% and applied at s0 values of 0.25 for TP-MS and 0.5 for PP-MS. For PP-MS volcano plot analysis, only proteins testing significantly enriched over the no DNA control (C_C) with FDR < 5% when tested using one-tailed Student’s t-testing, with permutation-based FDR control at 5% applied at an s0 value of 1 were considered for further analysis. All tested differences, *p*– values, and FDR-adjusted *q*-values are reported in supplementary tables.

### ZATT (ZNF451) tiling and analysis

sgRNA library targeting human ZATT (ZNF451) coding sequence was synthesized by GenTitan. For the generation of the sgRNA library, the sgRNAs were cloned into the base editor backbone Abe8e-Cas9-SpG lentiviral vector pRDA_479 (Addgene #179099) (Sangree et al. 2022) through Golden gate assembly with Esp3I and T7 DNA ligase. Briefly, the pool of oligonucleotides was amplified using NEBNext Ultra II Q5 Mastermix (New England BioLabs) with the primer set fwd 5’-AGGCACTTGCTCGTACGACG-3’, rev 5’-ATGTGGGCCCGGCACCTTAA-3’. The backbone was digested with Esp3I and gel extracted. All products were purified, before and after Golden gate cloning, using NucleoSpin Gel and PCR Clean-up (Macherey-Nagel). The sgRNA plasmid library was purified and concentrated via isopropanol precipitation. Endura ElectroCompetent Cells (Biosearch Technologies) were transformed to provide 1000 colonies per sgRNA. The cells were grown at 30°C for 15 hours on agar in the presence of 100 µg/ml carbenicillin before harvesting for plasmid purification (NucleoBond Xtra Maxi, Macherey-Nagel). To confirm library representation, a fraction of the purified sgRNA library was amplified, using i5 and i7 primers for multiplex indexing and NEBNext Ultra II Q5 Mastermix, gel extracted and sequenced using a NextSeq2000 (Illumina). For lentivirus production, HEK293T cells were transfected with the sgRNA plasmid library, psPAX2 (Addgene #12260) and pMD2.G (Addgene #12259) using Lipofectamine 3000 (Invitrogen) in Opti-MEM medium (Gibco) containing 5% FBS. The transfected cells were grown for 6 h before the media was replaced with fresh DMEM GlutaMax 10% FBS, 1% penicillin-streptomycin and 1% bovine serum albumin and then grown for an additional 42 h. Cells were then harvested, and the lentiviral supernatant passed through 0.45 μM filters for storage at –80°C. The screen was performed in RPE1-hTERT p53−/− cells. For lentiviral transduction, cells were incubated with the lentiviral supernatant at a multiplicity of infection between 0.2-0.3 and 8 mg/ml polybrene for 24 h. The transduced cells were then treated with 20 µg/ml puromycin for 24 h and re-seeded maintaining 20 µg/ml puromycin. Following 48 h of selection, cells were transferred to selection free media (T0, start-point). For the screen, cells were passaged every third day. Six days after selection, cells were split into an untreated, etoposide-treated (100 nM) or ICRF-193 treated (40 nM) fraction. The cells were passaged and treated accordingly for another 12 days (T18, endpoint). The drug doses correspond to the LD20, which were determined through a drug titration experiment in uninfected RPE1-hTERT p53−/− cells. The screen was performed in triplicates, with three separate transductions of the lentiviral library with a 1000-fold sgRNA coverage, which was maintained throughout the screen. Cell pellets were collected for each replicate at T0 and T18 for genomic DNA (gDNA) extraction. From the gRNA, the integrated sgRNAs were amplified using the primer set fwd 5’– GAGGGCCTATTTCCCATGATTC-3’, rev 5’-GTTGCGAAAAAGAACGTTCACGG-3’ and multiplexed with i5 and i7 barcodes in two successive PCR reactions using NEBNext Ultra II Q5 Master Mix. PCR2 products were purified by gel extraction and sequenced on an Illumina NextSeq2000. NGS sequencing data was processed by removing low abundance sgRNAs (counts < 30) and raw sequencing counts were normalized to log2 transcripts per million (log2tpm) per condition and replicate. The log2tpm values were compared for T0 vs etoposide-treated T18 and T0 vs ICRF-193 treated T18, and the fold change and p-value were calculated for each sgRNA using limma (Ritchie et al. 2015)

### Cell Culture

Cells were cultured in high-glucose DMEM with GlutaMAX Supplement and pyruvate (Gibco) supplemented with 10% FBS (Gibco) and 100 U per ml of penicillin-streptomycin (Gibco) at 37 °C with 5% CO2.

### Generation of cell lines

U2OS Flp-In T-REx cells were a kind gift from H. Piwnica-Worms. Four different gRNAs targeting different regions of ZATT (ZNF451) (5’-CCGAAGAGGAGGCCACACGT-3’, 5’– CCTCGGAGAACTATTTGCCG-3’, 5’-CTACCAGGGCATCTAAACCA-3’, 5’-TAAATCTTACAGTTGAGCGG-3’), were cloned into pSpCas9(BB)-2A-Puro (PX459) V2.0 (Addgene, 62988). sgRNA-containing plasmids were transfected into U2OS Flp-In T-REx cells using DharmaFECT kb (Horizon Discovert T-2006-01) transfection reagent according to the manufacturer’s protocol. After 24 h of incubation, transfected cells were selected with 1µg/mL puromycin for 48 h and plated sparsely to isolate single colonies. Single colonies were screened by immunoblot for a lack of ZATT protein and validated via sensitivity screen to etoposide using incucyte.

To generate pcDNA5/FRT/TO/EGFP-ZATT for ZATT KO reconstitution, codon-optimized hZATT cDNA was PCR-amplified and inserted 3’ to GFP in pcDNA5/FRT/TO/EGFP via KpnI and NotI sites. Derivative constructs expressing GFP-ZATT-*V134R and GFP-ZATT-*W137R were generated by the Q5 site-directed mutagenesis kit (New England BioLabs). ZATT KO U2OS Flp-In TREx cells stably expressing GFP-ZATT or point-mutation variants were generated by co-transfection with pOG44 (Invitrogen) and pcDNA5/FRT/TO/EGFP– ZATT plasmids (1:9 ratio) using FuGENE (Promega) according to manufacturer’s instructions. Selection for stable integration was performed with Hygromycin B (200 µg/mL, Invitrogen) and Blasticidin S (5 µg/mL, InvivoGen), and individual colonies were screened for expression with quantitative immunofluorescence and immunoblotting.

### Immunoblotting

To prepare whole-cell extracts, cells were lysed in EBC buffer (50 mM Tris-HCl; pH 7.5, 150 mM NaCl, 1 mM EDTA, 0.5% NP-40) for 30 min and sonicated for 30 s at 60% amplitude. Protein concentration was determined by the Bradford assay (Bio-Rad), and sample buffer (5% β-mercaptoethanol, 0.01% bromophenol blue) was added to all samples before boiling at 95 C for 5 min. Immunoblotting and development was performed as previously described (Poulsen et al. 2012).

### Sulforhodamine B (SRB) colorimetric assay

To assess cell growth via the SRB assay (PMID: 17406391), U2OS Flp-In TREx cells were grown in the presence of doxycycline (1 µg/mL, Clontech) 24 h before seeding them into 24 well plates at a density of 600 cells/well with indicated etoposide (ETP, Merck) concentrations and doxycycline. The cells were then left to grow for 12 d, whereafter they were washed with PBS, fixed with 10% trichloroacetic acid for 30 min at 4°C, washed twice with water and stained with 0.4% SRB in 0.1% acetic acid for 25 min RT. The plates were then washed 4 times with 0.1% acetic acid and left to dry overnight. Each well was resuspended in 100 µL 10 mM Tris-HCl pH 8.0 by incubating for 1 h at RT on a shaker. Absorbance was measured at A510 with an Epoch 2 microplate spectrophotometer (Agilent).

### CRISPR-Select cassette design

CRISPR-Select cassettes were designed as previous described (Niu et al. 2022). Briefly, we first designed gRNAs for SpCas9 targeting a PAM site close to the site of interest (max 10 bases away from the PAM site) on the gene. Then, a 45 bp long ssODN repair templates for knock-in were designed for both the mutant (MUT) and a synonymous WT control (WT’) at the same or nearby position as an internal control. As the knock-in site was within 10 bases to the PAM, this ensured that the gRNA seed region was destroyed after one round of knock-in. We also ensured that no splice site was targeted. If mutations on the ssODN templates were >10 bp away from the PAM sites, we included an additional synonymous mutation in the seed region to prevent gRNA rebinding. All oligos are listed in the table below.

### CRISPR-Select cassette delivery

gRNAs were transfected as a 75pM mixture of crRNA:tracrRNA duplexes purchased from Integrated DNA Technologies. In RPE1 p53^−/−^ Flag Cas9 cells, gRNAs were directly transfected as spCas9 is constitutively expressed in these cell lines. Following gRNA transfection, cells were transfected with a mixture of WT’ and MUT ssODNs at 10 pM. All transfections were performed using OptiMEM reduced serum media (Thermo Fisher Scientific, 31985062) and Dharmafect 1 transfection reagent (Horizon Discovery, T-2001– 03) as per manufacturer’s guidelines.

### CRISPR-Select^TIME^

CRISPR-Select^TIME^ was also performed as previously described (Niu et al. 2022). Briefly, 2 days after CRISPR-Select cassette delivery, an aliquot of the cells were collected to measure the ratio of MUT:WT’ ratio. The remaining cells were left in the dish and treated with 20% lethal doses of Etoposide (100 nM) for 10 days, with cells being replated in fresh media every 3 days. Finally, cells were collected again on day 12. Genomic DNA was extracted from cells using the Puregene cell kit (Qiagen) and 50 ng of genomic DNA was subjected to nested PCR, first using primer pairs (with overhangs for the second PCR) to amplify the locus of interest from the genome. Then, the second round of PCR was performed to add sample-specific barcodes and NGS adaptors for Illumina sequencing. These PCR products were purified and a 10 pM pooled amplicon sequencing library together with 20% PhiX was made and sequenced using the MiSeq Reagent Kit v2 (Illumina, MS-102-2002) in a MiSeq instrument (Illumina, SY-410-1003), according to manufacturer’s instructions. tructions. Sequencing depths ranged from 5,000 to 50,000 reads per sample. NGS data were analyzed by the CRISPResso2 online tool using default settings (Clement et al. 2019).

Oligos used for CRISPR-Select^TIME^

### gRNAs

TOP2A-E979G

GACTGAAGAAAAACTGGCAG

TOP2A-V988Y

AGTTGCATGTGAGACTAGTT

### ssODN repair templates

TOP2A-E979G

MUT:ACCACTGTGAAATTTGTTGTGAAGATGACTGAAGAAAAACTGGCAGGCGCAGAGAGAGTTGGACTACACAAAGTCTTCAAACTCCAAACT

WT’:ACCACTGTGAAATTTGTTGTGAAGATGACTGAAGAAAAACTGGCAGAAGCAGAGAGAGTTGGACTACACAAAGTCTTCAAACTCCAAACT

TOP2A-V988Y

MUT:ACTGAAGAAAAACTGGCAGAGGCAGAGAGAGTTGGACTACACAAATACTTCAAACTCCAAACCAGTCTCACATGCAACTCTATGGTATGT

WT’:ACTGAAGAAAAACTGGCAGAGGCAGAGAGAGTTGGACTACACAAAGTATTCAAACTCCAAACCAGTCTCACATGCAACTCTATGGTATGT

### PCR 1 – primers

Forward:ACACTCTTTCCCTACACGACGCTCTTCCGATCTTGGCACCGAGAAGACAC

CTCCT

Reverse:TGACTGGAGTTCAGACGTGTGCTCTTCCGATCTTGAAACGTGTACATCTCA

CAAAACA

### Data Availability

The mass spectrometry proteomics data have been deposited to the ProteomeXchange Consortium via the PRIDE (Perez-Riverol et al. 2022) partner repository with the dataset identifier PXD070044.

### Reproducibility

A minimum of two independent experiments has been conducted for each experimental result shown in this manuscript.

## Author Contributions

X.L. performed all experiments unless stated otherwise; A.Z. developed the TOP2cc release assay (Fig.5G-H, S5B-C) and assisted with protein purification of ZATT proteins; E.K., S.A.G., and S.M. performed the CRISPR base editor screen (Fig. 4A-D); N.E.D. designed the guide RNA library of the CRISPR base editor screen and analyzed the sequencing data with assistance of S.A.G.; S.A.G. and S.M. performed the CRISPR Select experiment with assistance of S.W. under the supervision of C.S.S. (Fig.7D-E, S7B); P.F. purified some TOP2a and ZATT variants and performed the depletion addback experiments in Fig.6H and S6D, relaxation assay (Fig.S5A); I.H. acquired and analyzed mass spectrometry samples under the supervision of M.L.N.; C.S.C. assisted with the mass spectrometry analysis; E.K. and A.I. generated the human ZATT cell lines; A.I. performed the cell growth assays under the supervision of N.M. (Fig.7A-C, S7A); S.K. established the TrimAway assay in the lab; C.G. designed and developed the ALFA-tag pulldown assay under the supervision of T.C.R.M.; N.P. performed the alignment analysis (Fig.6E, S6I); T.C.R.M. assisted with AF3 predictions; A.G.L. participated in the initial phases of this project and TOP2a purification; X.L. and J.P.D. designed and analyzed the experiments, and prepared the manuscript with feedback from all authors.

## Acknowledgments

We thank Jiri Lukas, Johannes Walter, and all members of the Duxin laboratory for critical feedback on the manuscript. We also thank Yoshi Azuma for sharing valuable reagents. We are grateful to Sara Ambjørn, Bob Meeusen, Emil Hertz, and Jakob Nilsson for their assistance in establishing CRISPR genome-editing approaches in the laboratory. This project has received funding from Cancer Research UK Senior Cancer Research Fellowship (C68484/A28159), Novo Nordisk Foundation (NNF23OC0085973, NNF22OC0074140, NNF24SA0098829, and NNF22SA00782290), European Research Council under the European Union’s Horizon 2020 research and innovation program. This project was Funded by the European Union. Views and opinions expressed are however those of the author(s) only and do not necessarily reflect those of the European Union or European Research Executive Agency. Neither the European Union nor the granting authority can be held responsible for them.

## Supplemental Figure Legends

**Figure S1.**
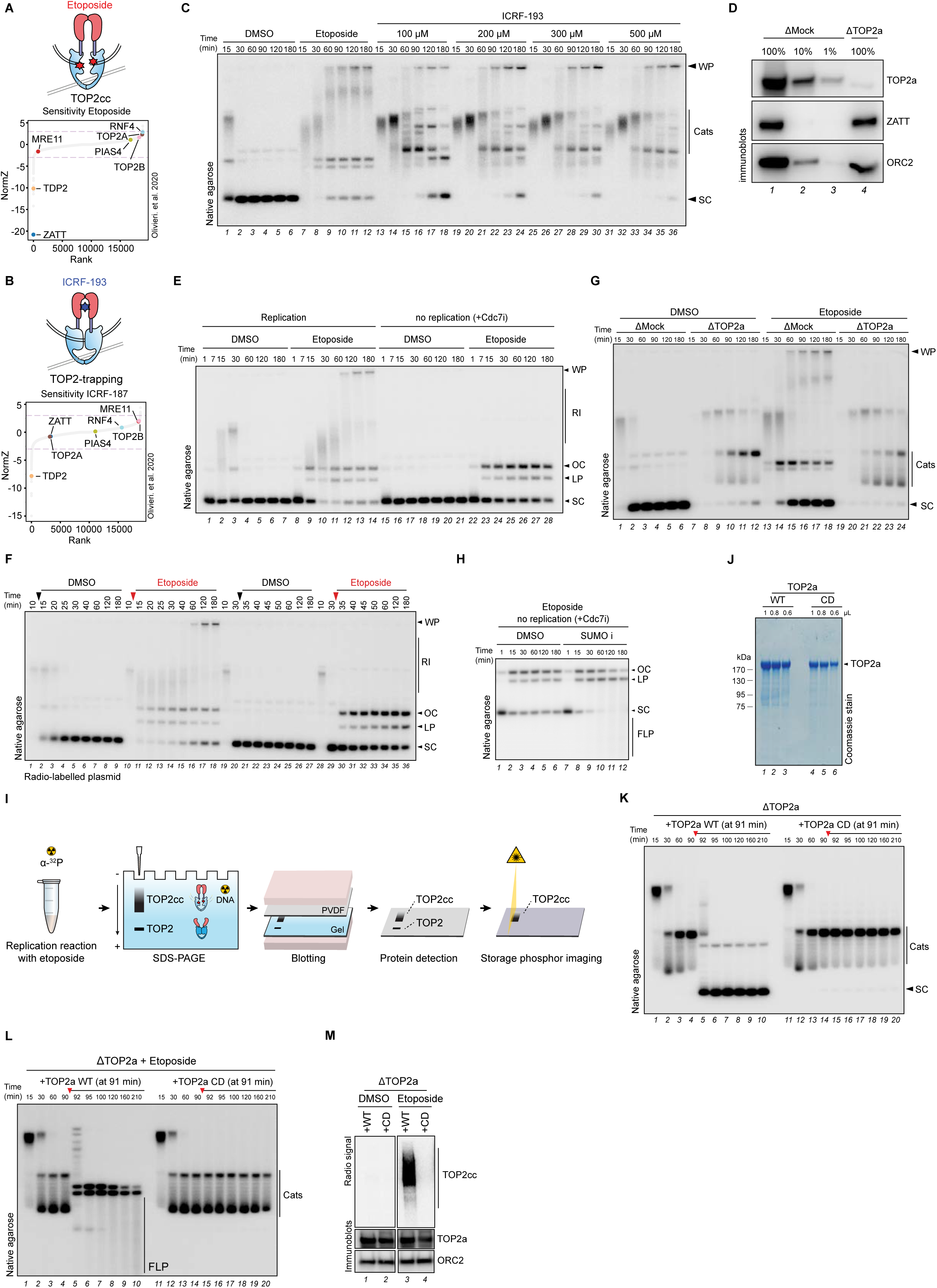
**A**) Normalized gene-level Z-scores for a genome-wide sensitivity dropout screen to etoposide in RPE p53 KO cells (Olivieri et al. 2020). **B**) Normalized gene-level Z-scores for genome-wide sensitivity dropout screen to ICRF-187 RPE p53 KO cells (Olivieri et al. 2020). **C**) pBS was replicated in egg extracts in the presence of DMSO, etoposide (100 µM), or the indicated concentrations of ICRF-193. Samples were analyzed as in Fig.1C. Cats, catenanes. **D**) Mock– or TOP2a-depleted extracts were immunoblotted with the indicated antibodies. **E**) Pre-labelled pBS was replicated in egg extract in the presence of DMSO, etoposide (100 µM). Cdc7 inhibitor was supplemented to reaction as indicated. Samples were analyzed as in Fig.1C. Note that the WP only appeared in the presence of etoposide and replication. **F**) pBS was replicated for 10 or 30 minutes prior to the addition of either DMSO (black arrow) or etoposide (100 µM, red arrow). Samples were analyzed as in Fig.1C. Note that WPs appear only when etoposide is added during DNA replication at 10 minutes. **G**) pBS was replicated in mock– or TOP2a-depleted extracts in the presence or absence of etoposide (100 µM). Samples were analyzed as in Fig.1C. **H**) Pre-labelled pBS was incubated in egg extract in the presence of etoposide with or without SUMO inhibitor. Cdc7 inhibitor was supplemented to block DNA replication. Samples were analyzed as in Fig.1E. **I**) Illustration of drag-on blot workflow. Replication reaction with [α-32P] dATP is analyzed on SDS-PAGE, followed by blotting onto PVDF membrane. Protein is detected by immunoblot with specific antibodies and cross-linked DNA is analyzed by storage phosphor imaging. **J**) Purified recombinant Xenopus TOP2a wildtype (WT) or catalytic dead (CD) were resolved via SDS-PAGE and stained with Coomassie blue. **K**) pBS was replicated in TOP2a-depleted extracts. At 91 minutes either TOP2a WT or CD were supplemented to the reaction where indicated (red arrow). Samples were analyzed as in Fig.1C. **L**) Same as (K) but in the presence of etoposide (100 µM). **M**) Samples from (K) and (L) were analyzed by drag-on blot (radio signal) and immunoblot.

**Figure S2.**
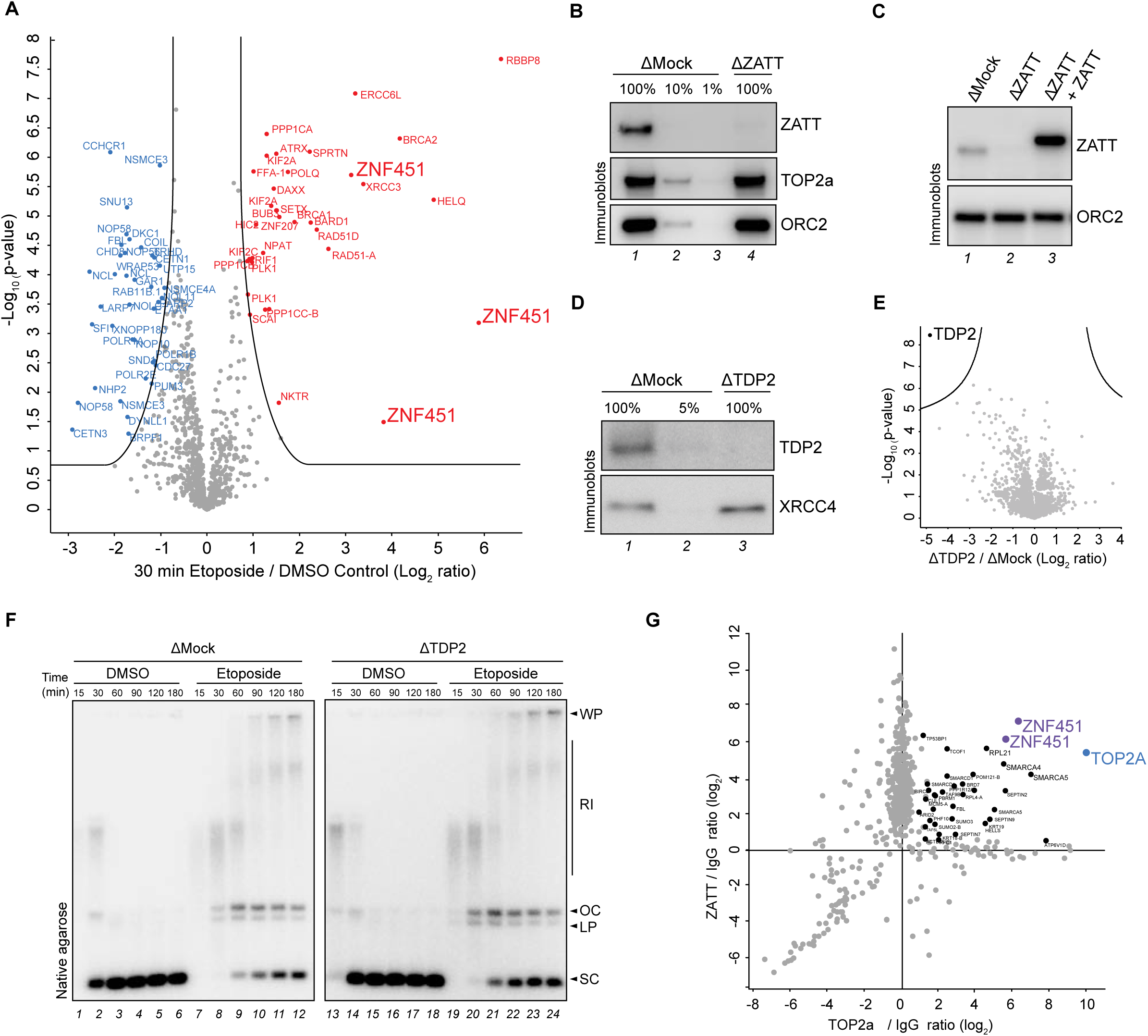
**A**) PP-MS volcano plot analysis comparing protein enrichment to plasmids between the etoposide and DMSO (control) reaction at 30 minutes. The volcano plot shows the difference in abundance of proteins between the two sample conditions (x– axis), plotted against the p-value resulting from two-tailed Student’s two-sample t-testing (y-axis). Proteins significantly down– or up-regulated (cutoff line represents permutation– based FDR<5%) are represented in blue or red, respectively. n=4. Full results reported in Table S1. **B**) Extracts that were either mock– or ZATT– depleted were immunoblotted with the indicated antibodies. **C**) Mock– or ZATT-depleted extracts were supplemented with wildtype (WT) recombinant ZATT where indicated and immunoblotted with the indicated antibodies. **D**) Mock– or TDP2-depleted extracts were immunoblotted with the indicated antibodies. XRCC4 was used as a loading control. **E**) Total proteome MS analysis of mock– versus TDP2-depleted egg extracts. The volcano plot shows the difference in abundance of proteins between the mock– and TDP2-depleted samples (x-axis), plotted against the p-value resulting from two-tailed Student’s t-testing with permutation-based FDR-adjusted p-value of < 0.05 (y-axis). n=4 biochemical replicates. Full results are reported in Table S2. **F**) pBS was replicated in mock– or TDP2-depleted extracts in the presence or absence of etoposide (100 µM). Samples were analyzed as in Fig.1C. **G**) pBS was replicated in the presence of etoposide (100 µM) and samples collected at 30 min and immunoprecipitated with either IgG, ZATT, or TOP2a antibody. Plot shows the fold enrichment of TOP2a over IgG (x-axis) over the fold enrichment of ZATT over IgG (y– axis). Labelled dots are proteins significantly enriched with TOP2a and ZATT immunoprecipitation when evaluated by two-tailed Student’s t-testing, with permutation– based FDR control at 5% and applied at s0 values of 0.25. n=4 biochemical replicates. Full results are reported in Table S3.

**Figure S3.**
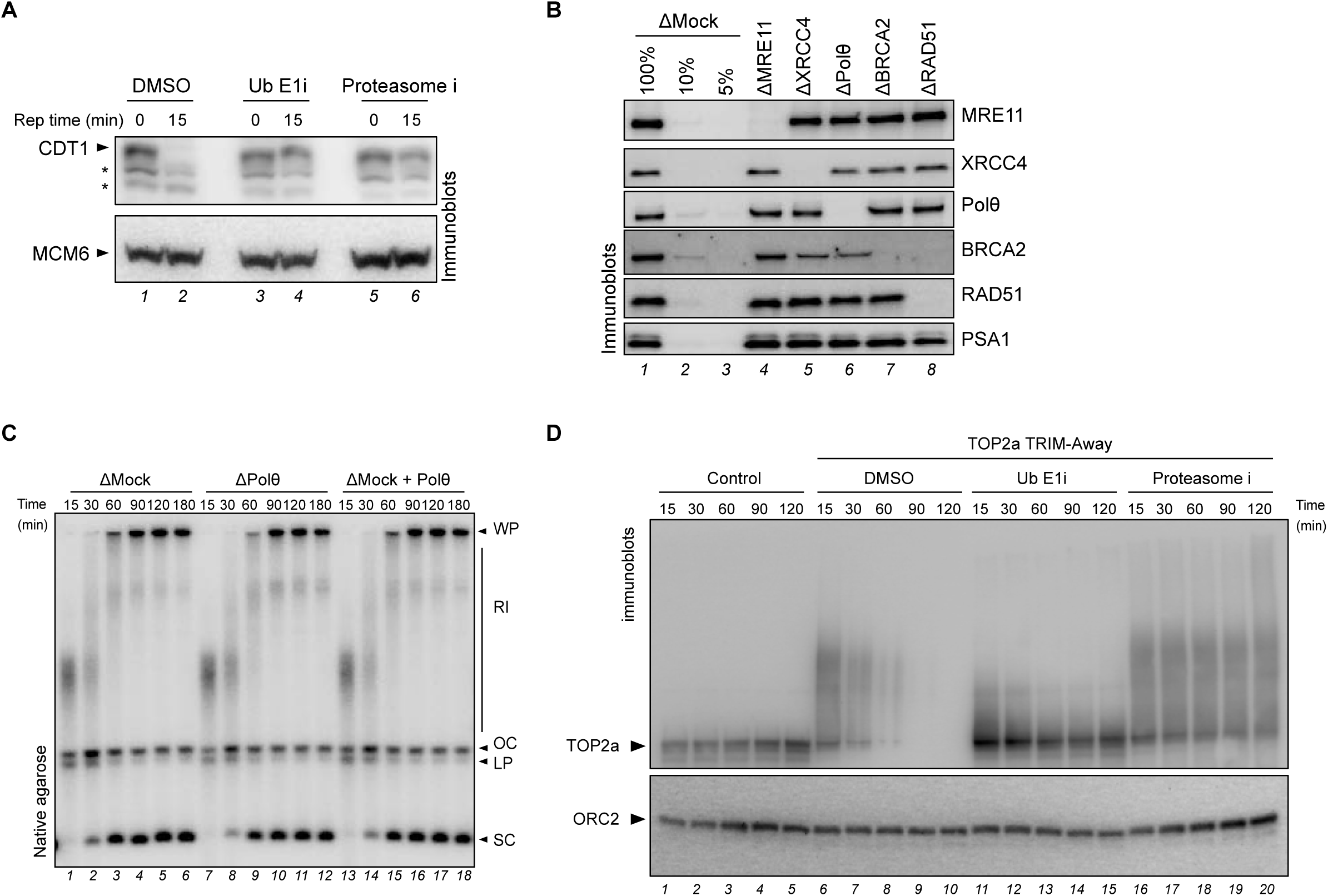
**A**) Functional validation of Ub E1 inhibitor (Ub E1i, MLN7243) and proteasome inhibitor (Proteasome i, MG262) by monitoring CDT1 destruction during DNA replication. Note that CDT1 destruction was hindered with both Ub E1 and proteasome inhibition. **B**) Mock-, MRE11-, XRCC4-, Polθ-, BRCA2-, or RAD51-depleted extracts were immunoblotted with the indicated antibodies. PSA1 was used as loading control. **C**) pBS was replicated in either mock– or Polθ-depleted extracts. Mock-depleted extracts were also supplemented with Polθ inhibitor (Polθ i) where indicated. Samples were analyzed as in Fig.1C. **D**) Immunoblot of TOP2a degradation by TRIM-Away in extract supplemented with Ub E1i or proteasome i. Note that TRIM-Away induced TOP2a degradation is blocked by Ub E1 or proteasome inhibition.

**Figure S4.**
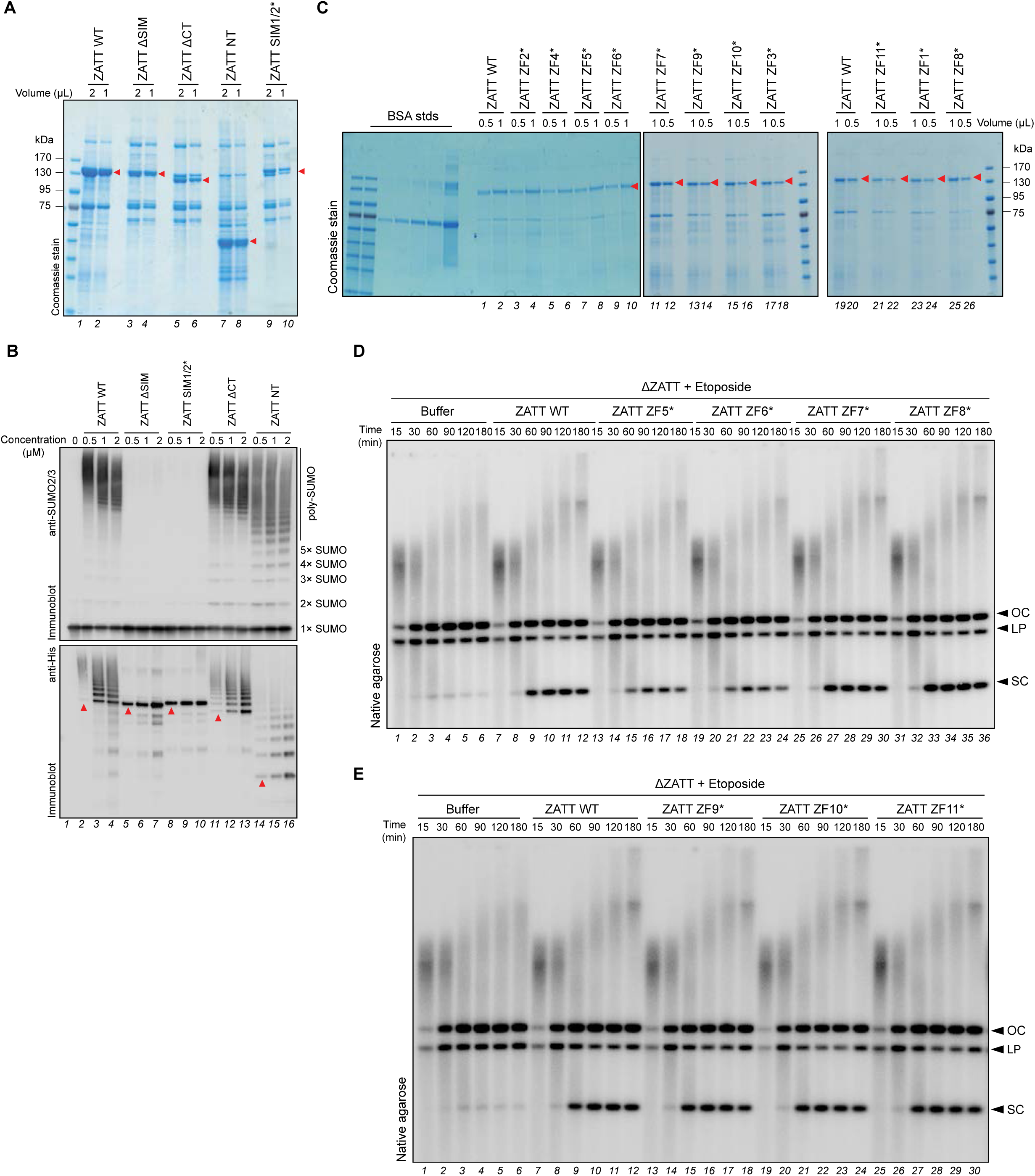
**A**) ZATT wildtype (WT) and the indicated ZATT mutant proteins (red arrows) were purified and analyzed by SDS-PAGE and Coomassie staining. **B**) ZATT wildtype (WT) and indicated ZATT mutants were tested for their ability to catalyze poly– SUMOylation *in vitro*. Recombinant WT and mutant ZATT proteins were incubated at the indicated concentrations with SAE1, UBA2, UBC9 and SUMO2, and analyzed by immunoblotting with the indicated antibodies. Red arrows indicate corresponding recombinant 6×His-tagged ZATT. **C**) The indicated ZATT zinc finger mutants (red arrows) were purified and analyzed by SDS-PAGE and Coomassie staining. **D**) pBS was replicated in ZATT-depleted extracts in the presence of etoposide (100 µM). Extracts were supplemented with buffer or the indicated mutants and analyzed as in Fig.1C. **E**) Same as (D) but with different ZATT mutants.

**Figure S5.**
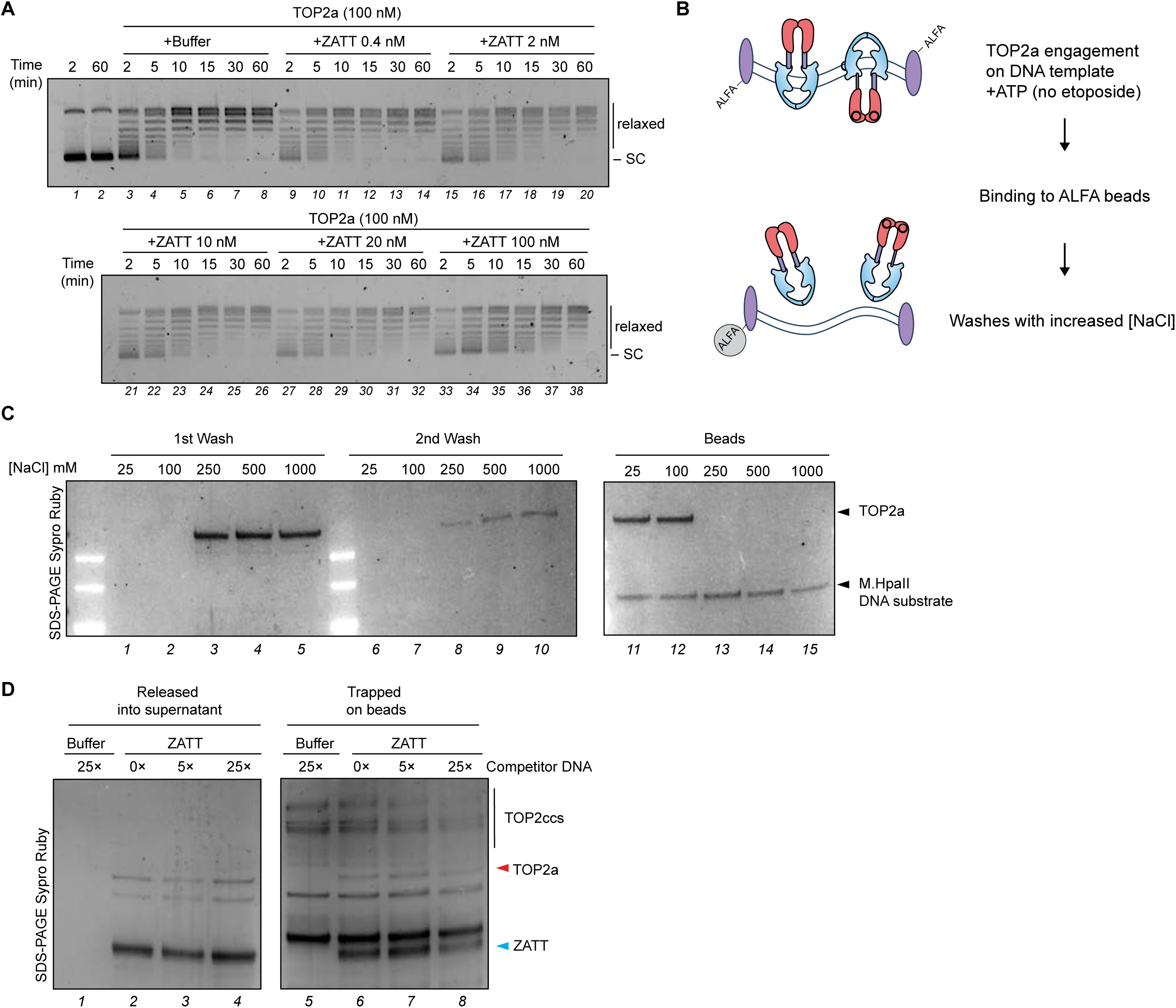
**A**) Relaxation of undamaged pBS plasmid by TOP2a (100 µM) supplemented with ZATT at indicated concentration. **B**) Illustration of TOP2a covalent trapping assay. The stringent wash is performed to remove non-covalent TOP2a-DNA interactions. **C**) DNA-TOP2a pulldown performed in the absence of etoposide. Non-covalent TOP2a-DNA complexes were washed with the indicated NaCl concentrations and supernatants (1^st^ and 2^nd^ wash), and beads were resolved via SDS-PAGE and stained with SYPRO Ruby. Results show that high concentrations of NaCl (> 250 mM) release non-covalently bound TOP2a from DNA. **D**) TOP2cc DNA-bead complexes were incubated with buffer or recombinant ZATT for 30 minutes in the absence or presence of competitor DNA. 0×, 5× and 25× corresponds to no addition of competitor DNA, competitor DNA in 5 times, or 25 times the molar concentration of the linear DNA substrate added to the release reactions. Supernatant (released) and beads (trapped) were recovered and analyzed by SDS-PAGE gel followed by SYPRO Ruby staining. Red arrow indicates TOP2a, blue arrow indicates ZATT.

**Figure S6.**
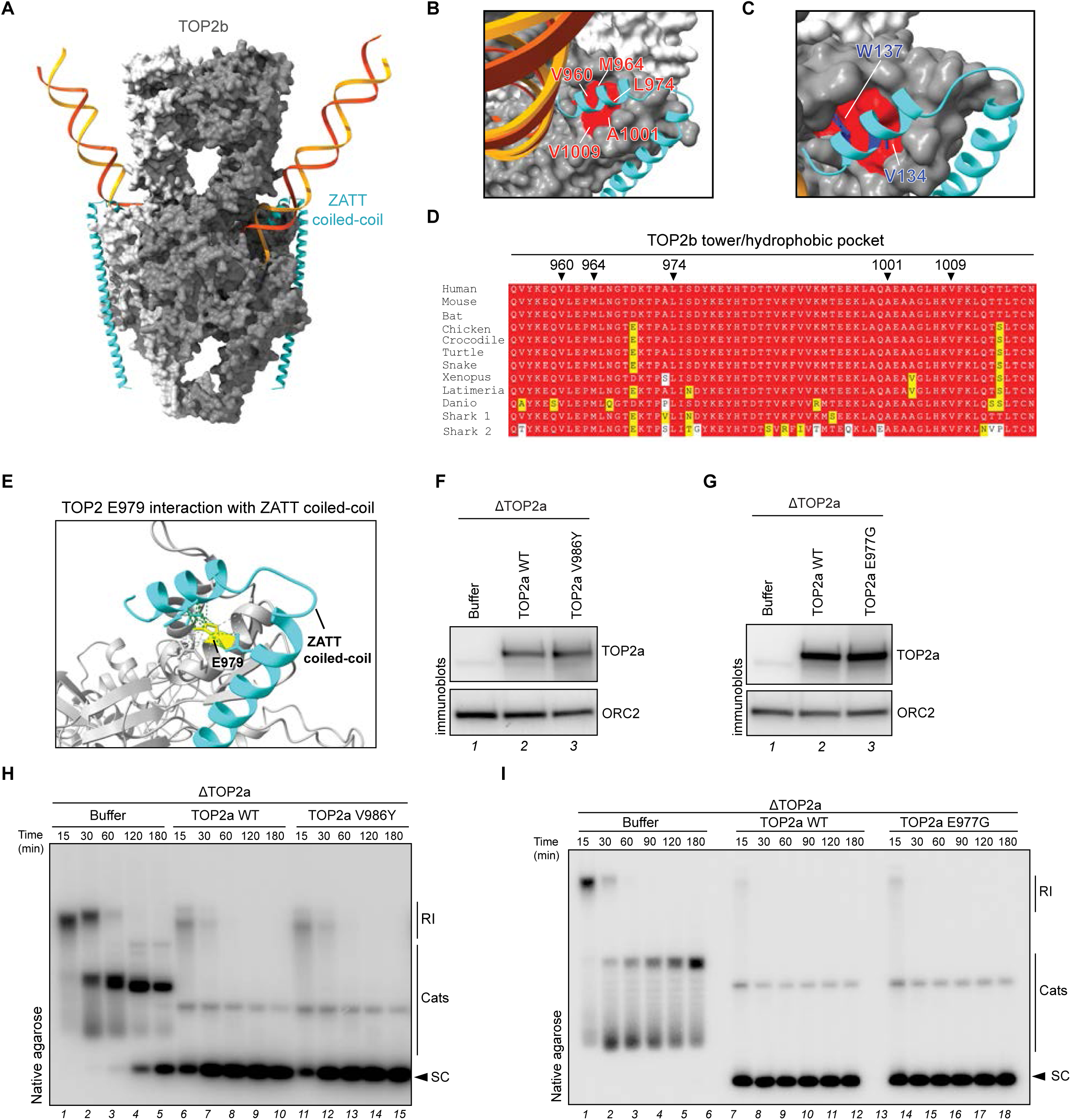
**A**) AlphaFold3 predicted model of human TOP2bcc and the ZATT coil-coiled domain. **B**) Close up view of (A) highlighting the human TOP2b hydrophobic pocket residues (in red). **C**) Close up of (A) highlighting human ZATT W137 and V134 (in blue). **D**) Multiple sequence alignments of TOP2B from different species. Conserved residues are highlighted in red, conservative substitutions in yellow, and non-conservative substitutions in white. TOP2b residues located within the hydrophobic pocket are indicated with arrows. **E**) Close up view of AlphaFold3 predicted model of human TOP2a dimer and ZATT coil-coiled domain illustrating the contacts made by TOP2a E979 (yellow) with the ZATT hook domain (cyan). **F**) Egg extracts depleted of TOP2a and supplemented with buffer, *Xenopus* TOP2a WT, or TOP2a V986Y (corresponding to V988 in human) were immunoblotted with indicated antibodies. **G**) Egg extracts depleted of TOP2a and supplemented with buffer, *Xenopus* TOP2a WT, or TOP2a E977G (corresponding to E979 in human) were immunoblotted with indicated antibodies. **H**) Extracts from (F) were used to replicate pBS. Reaction samples were analyzed as in Fig.1C. **I**) pBS was replicated in extracts from (G). Reaction samples were analyzed as in Fig.1C.

**Figure S7.**
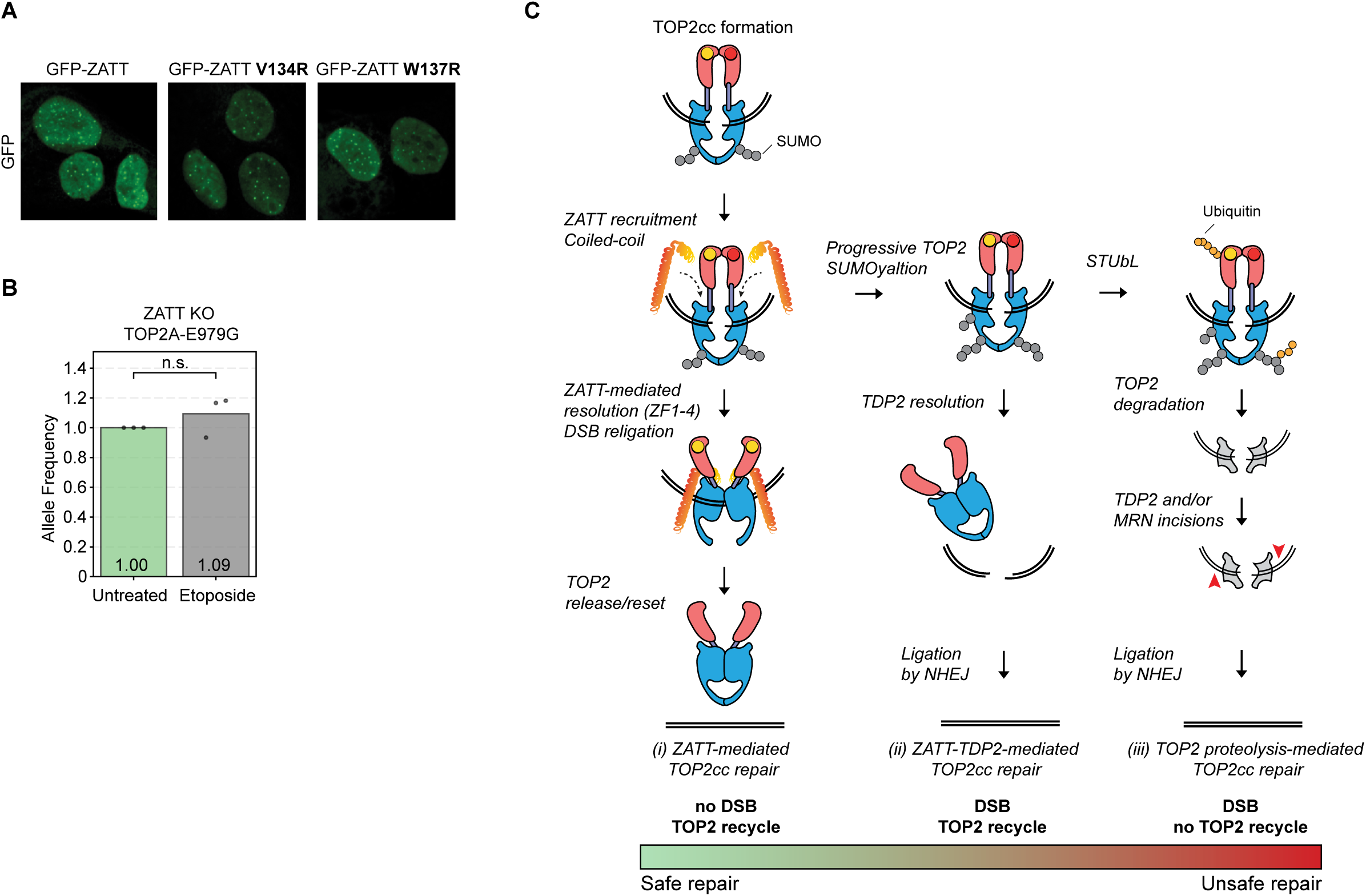
**A**) Representative images of GFP fluorescence of the indicated U2OS cell lines. **B**) CRISPR-Select analysis of TOP2a-E977G in ZATT KO U2OS cells treated or untreated with etoposide for 12 days. Allele frequency was measured by NGS, normalized to untreated value. Means of 3 independent biological replicates are shown, n.s. denotes no significance by Student’s t-test. **C**) Illustration of the different models of TOP2cc repair based on this work (i) and previous work (ii and iii).

## Table legends

**Table S1**. PP-MS analysis related to Fig.1H and Fig.S2B. C, DMSO control reaction; Et, etoposide reaction; Et-Si etoposide and SUMOi reaction.

**Table S2**. Whole proteome MS analysis related to Fig.S2F. Control (C), mock-depleted extracts; ΔTDP2, TDP2-depleted extracts.

**Table S3**. IP-MS analysis related to Fig.S2H. IgG, control immunoprecipitate; ZATT, ZATT immunoprecipitate; TOP2a, TOP2a immunoprecipitate.

**Table S4**. ZATT base editor tiling screen results and analysis related to Fig.3. The significance and fold change were derived from limma (see methods).

**Table S5**. AlphaFold 3 predictions related to Fig.4E-H and Fig.S6A-C. Note that in Table S5, the predicted interaction with ZF1 (in grey) is incompatible with the TOP2a dimer structure. ZATT coiled-coil (C-C) predicted interactions with TOP2a and TOP2b are highlighted in green.

**Table S6**. CRISPR-Select results related to Fig.4O and Fig S5O.

**Table S7**. PCR primers used to make TOP2a and ZATT variants.

